# Seed fossil record of Solanaceae revisited

**DOI:** 10.1101/2025.07.03.662944

**Authors:** Rocío Deanna, Aleksej Vladimirovič Hvalj, Edoardo Martinetto, Sandra Knapp, Eva-Maria Sadowski, Steven Manchester, Abel Campos, Vincent Fernandez, Gloria E. Barboza, Hervé Sauquet, Ellen Dean, Tiina Särkinen, Franco Chiarini, Gabriel Bernardello, Stacey D. Smith

## Abstract

The fossil record for Solanaceae has a complex taxonomic history, with many species originally described in the family being subsequently shown to belong in other plant families. In this work, we present an in-depth analysis of the nightshade seed fossil record, which corresponds to the largest amount of fossil material for the family despite having relatively few described taxa. We re-examined 110 fossil seeds previously assigned either to one of the six taxa or unplaced within the family. All fossils were imaged and measured for 14 discrete and continuous characters, and a subset of 22 well-preserved specimens was additionally subjected to micro-CT scanning. In parallel, we scored the same set of characters for seeds from 354 extant taxa of Solanaceae. We carried out a multivariate analysis of the combined extinct and extant dataset to visualize the placement of fossil seeds within Solanaceae seed morphospace. This analysis revealed a wide spread of fossils across morphospace, including a variety of shapes, sizes and testal wall features. For one fossil taxon (*Hyoscyosperma daturoides* Deanna & S.D.Sm. gen. et sp. nov.), the micro-CT images confirmed the presence of an elaiosome, otherwise only known in the family from the genus *Datura* L. Based on these analyses, we re-circumscribe three species and describe six new taxa (four in new genera), doubling the number of fossil seed species in the family. The broad geographic and stratigraphic distribution of these diverse taxa, from Eocene North America and Europe to Pleistocene Eurasia, support the notion that the Solanaceae is likely a family of a late Cretaceous origin that attained nearly a cosmopolitan distribution by the Eocene.

## INTRODUCTION

The nightshade family, Solanaceae, comprising roughly 3,000 species and 99 genera, is known for its economic and ecological importance (Knapp & al., 2004) as well as its diverse array of specialized metabolites (Eich, 2008), which together drive ongoing research in genomics, plant breeding and biodiversity science (Pozcai & al., 2022). Nevertheless, understanding the biogeographical context and timeline of the nightshade radiation has been hindered by the lack of information from the fossil record that has received relatively little focused attention until recently. Martinez-Millan (2010) provided the first review of fossils described in the family and listed 11 taxa, based on seeds, leaves, pollen, and flowers. However, a subsequent revision (Millan & Crepet, 2014) concluded that none of these fossils presented enough characters to be confidently assigned to Solanaceae, suggesting that one genus of fossil flowers (*Solanites* Saporta) instead belonged to Rhamnaceae. The nightshade fossil record also includes two wood fossils (Page, 1980; Franco & Brea, 2008), the latter of which has been suggested to have affinities to *Solanum* L. (Franco & Brea, 2023). Most recently, four new Solanaceae taxa based on fossil berries were described from South America (*Physalis infinemundi* Wilf*, Physalis hunickenii* Deanna, Wilf & Gandolfo*, Lycianthoides calycina* Deanna & Manchester*, Eophysaloides inflata* Martínez C.M. & Deanna; Wilf & al., 2017; Deanna & al., 2020; Deanna & al., 2023). Unlike other fossils of Solanaceae, these fossilized berries are rich in informative characters (e.g., calyx venation pattern, pedicel insertion), which have allowed them to be placed in tribes or even genera within the family (Deanna & al., 2023).

The Solanaceae seed fossil record presents a large reservoir of potential new information about the evolutionary history of the family. Although there are only six described seed fossil taxa, four of these are represented by multiple specimens, and there is also a large pool of fossil seeds that have been assigned to extant taxa (e.g., Martinetto, 1995; Basilici & al., 1997; Cavallo & Martinetto, 2001; Niccolini & al. 2022; reviewed in Särkinen & al., 2013, 2018). Särkinen & al. (2018) analyzed three of the extinct seed taxa and determined that the best preserved (*Cantisolanum daturoides* Reid & M.Chandler) is likely a commelinid monocot. The two remaining taxa, *Solanispermum reniforme* M.Chandler and *Solanum arnense* M.Chandler, likely belong to Solanaceae (Särkinen & al., 2018) but have only enough evidence to be placed in the large Solanoideae clade or within the crown group of the family (Särkinen & al., 2013; De-Silva & al., 2017). The other three fossil seed taxa (*Physalis pliocenica* Szafer, Szafer, 1946; *Solanum foveolatum* Negru, Negru, 1986; *Nephrosemen* Manchester, Manchester, 1994) have not been analyzed in the context of the broader fossil record of the family. In addition, some of these fossil seeds historically assigned to extant species may represent extinct taxa rather than members of living lineages.

Seeds in the Solanaceae share features that help to distinguish them from those in other plant families, but also present informative variation that facilitates assignment to particular clades within the family. All solanaceous seeds are campylotropous, i.e., they have a bent longitudinal axis, where the chalazal and micropylar ends meet close to the hilum. Most of the family (clade Solaneae sensu Särkinen & al., 2013) additionally exhibits flattened seeds generally sized between 1.5-2.5 mm and reniform in shape with curved to circinate embryos (Särkinen & al., 2018). Members of other solanaceous clades, such as the subfamilies Nicotianoideae and Petunioideae (Särkinen & al., 2013), have generally smaller (< 1 mm) and not flattened (e.g., angular, globose) seeds with straight embryos, but these seeds are still campylotropous in that the hilum is positioned sublaterally (Hunziker, 2001; Fig. 5 in Särkinen & al., 2018). Some seed traits are rare but highly diagnostic, such as the elaiosomes of *Datura* L. and the asymmetric wings of *Sessea* Ruiz & Pav. (Hunziker, 2001). Seeds in the family also present notable continuous size variation, e.g., ranging from tiny seeds (<1 mm in Nicotianoideae) to the size of a small mango seed (ca. 30 mm long in *Duckeodendron* Kuhlm.).

Here we apply state-of-the-art imaging methods along with morphometric analyses to revise the Solanaceae seed fossil record. We imaged 110 seed fossil specimens from collections and literature, and we measured 14 discrete and continuous characters to capture their variation in size, shape, ornamentation, and other features. For 22 of these seeds with sufficient preservation, we also conducted X-ray micro-Computed Tomography (micro-CT) to examine both the internal and external structure, as this has been instrumental in previous seed fossil analyses (Särkinen & al., 2018). In order to place these fossil specimens in the context of the morphological diversity of the family (Gandolfo & al., 2008; Parham & al., 2012), we scored additional 354 extant taxa for the same sets of seed characters. Our results reveal that the fossil seeds span much of the morphospace of extant seeds, including both medium-sized flattened and small angular forms as well as taxa with and without elaiosomes. In total, we identify 17 distinct morphological clusters of fossil seeds, 10 of which include specimens scored directly from fossils (as opposed to only the literature). Focusing on these 10 clusters, we re-circumscribe three fossil species, make one new combination, and describe four new fossil genera and six new fossil species. These efforts double the described diversity of Solanaceae fossil seeds and, as we discuss, provide new insights into the evolutionary and biogeographic history of the family.

## MATERIAL AND METHODS

### Revision of seed fossils of Solanaceae

Carpological nightshade collections by Pavel Ivanovich Dorofeev, Hans Dieter Mai, Marjorie Chandler, Eleanor Reid, Steven Manchester, and Vadim Petrovich Nikitin were studied at North American, European and Russian museums. In Italy, the CENOFITA collection, managed by the Regional Museum of Natural Sciences of Turin (acronym MRSN-P/345-CCN), was studied. All fossilized seeds were analyzed and photographed using a dissecting microscope at different institutions (Natural History Museum, London, UK; Museum für Naturkunde Berlin, Germany; Florida Museum of Natural History, Gainesville, US; Komarov Botanical Institute of the Russian Academy of Sciences, Saint Petersburg, Russia; Universitá degli Studi di Torino, Torino, Italy; Naturalis Biodiversity Center, Leiden, The Netherlands). All previous descriptions and synopses were analyzed (e.g., Chandler, 1957; Palamarev, 1970; Dorofeev, 1963; Mai, 2001; Velichkevich & Zasfawniak, 2003).

### Classification system

Circumscription of Solanaceae herein follows Särkinen & al. (2013). The family includes 99 genera and almost 3,000 species arranged in six subfamilies (Hunziker, 2001). We followed World Flora Online (WFO; https://www.worldfloraonline.org/) for accepted names of all extant taxa. We checked generic and specific names of new taxa against the International Plant Name Index (https://ifpni.org/index.htm) and the International Fossil Plant Names Index (https://ifpni.org/).

### Scoring of extant and fossil seeds

In order to assess the morphological diversity and taxonomic affinities of the 110 fossil seed specimens, we assembled a dataset comprising both continuous and discrete characters for extinct and extant taxa. For 354 living species and 110 fossils, we scored two continuous traits (seed length and width) and seven discrete traits (e.g., compression, hilum position, embryo shape; see Table S2 for complete list and coding scheme). This large data set included at least one species for each of the 99 genera, except for *Henoonia* Griseb., *Poortmania* Drake, and *Mellissia* Hook.f., for which we were unable to obtain material. Of the 110 fossils, 95 were scored from collections (i.e., physical material) and 15 were scored from literature alone (i.e., protologues). Seed traits were scored from living material, herbarium specimens, micro-CT, and taxonomic literature (Tables S1, S3, S4). For 159 of the extant species and 95 of the fossil specimens, we also imaged the seeds in a dissecting microscope. Image stacks of fossils were manually taken with digital cameras that were installed on the microscopes at different institutions (see above). For the 159 extant taxa, seeds were collected from herbaria and carpological collections (A, BM, BR, COLO, CORD, DAV, L, LE, MO, NY) and the image stacks were automatically taken with digital cameras installed on stereomicroscopes either at CU Boulder or at the NHM. By applying the software package of HeliconFocus Pro, the stacks were then digitally merged into photomicrographic composites, available as part of the OSF repository associated with this publication (link will be provided after revisions). These 254 images were used to measure or calculate five additional continuous traits: area, perimeter, ratio area/perimeter, and roundness of exotestal cells at the center of the seed surface, and seed total area. We measured area, perimeter and roundness of three exotestal cells at the center of the seed surface, and calculated an average using Image J (Schneider & al., 2012). In total, between the initial scoring and the analysis of microscope images, we gathered data for 14 morphological characters (seven continuous and seven discrete). All data were recorded in the PROTEUS database (Sauquet & al., 2017) linking each record to an explicit source or voucher (Tables S1, S3, S4).

### X-ray computed tomography

In total, 22 seed fossils were scanned using X-ray computed tomography (Table S5) to evaluate pyritization and infer internal preservation, including two discrete characters: embryo shape and presence of elaiosomes. In addition, 14 extant seeds were scanned using the same method to compare the internal structure with that of the fossils. These included representatives of all genera previously associated with seed fossils (e.g., *Physalis* L.*, Scopolia* Jacq.*, Solanum* L.) as well as additional taxa selected to capture the broader morphological diversity of Solanaceae seeds (e.g., *Browallia* L.*, Nicotiana* L.*, Schizanthus* Ruiz & Pav.; Table S5). A high-resolution Zeiss Versa 520 micro-CT system, which combines classic X-ray cone beam magnification with optical lenses (0.4X, 4X, 20X and 40X) to gain additional magnification, was used to acquire the scans with 1600 projections over 360°. Seeds were removed from their glass cylinders and placed into a small piece of floral foam to allow higher X-ray penetration and image resolution. The scans were performed using a transmission tungsten target, applying the following settings: voltage = 50-100 kV, current = 80-90 µA, power = 4-9 W, and exposure time = 2-12 s. The complete list of parameters is detailed in Table S5. The visualization of the images and cutting of the volume files into image stacks were done using the software Volume Graphics Studio Max 3.4 (Volume Graphics, Heidelberg, Germany).

### Morphospace analysis

We used our dataset of 14 morphological traits (Table S2) scored for 464 specimens (Tables S1, S3) to perform a cluster analysis to compare the fossil seeds with those of extant Solanaceae. We applied a non-metric multidimensional scaling (NMDS) analysis to assess the similarity of the fossils to seeds of extant taxa, identifying clusters according to morphology as it was previously done with Solanaceae fruit macrofossils in Deanna & al. (2023). We carried out NMDS with the metaMDS function and Gower’s Distance as the dissimilarity metric in the VEGAN package in R (Oksanen & al., 2013). All characters were treated as symmetric except embryo shape, which was considered ordered (straight to curved to coiled). We also ran a stress analysis to test how many dimensions are needed to plot the data (Fig. S1). Results were plotted using the function ggplot from the R GGPLOT2 package (Wickham & al., 2016).

## RESULTS

### Micro-CT of fossil and extant Solanaceae seeds

We recovered valuable internal structural data from three fossil specimens out of the total 22 fossil seeds and 14 extant seeds analyzed using high-resolution micro-CT. These exceptionally preserved fossils revealed key anatomical features, including the presence of an elaiosome attached to the seed surface (BIN K453/133; **Fig. 1A-B**) and the curvature and position of the embryo (e.g., curved in MB.Pb.1993/6802 (**Fig. 1C**) vs. coiled in BIN K453/175; **Fig. 1D**). The visibility of these internal traits not only confirms the campylotropous seed type of Solanaceae but also provides insight into seed anatomy beyond external morphology, allowing for more confident placement of fossil taxa within the family.

**Figure 1.**
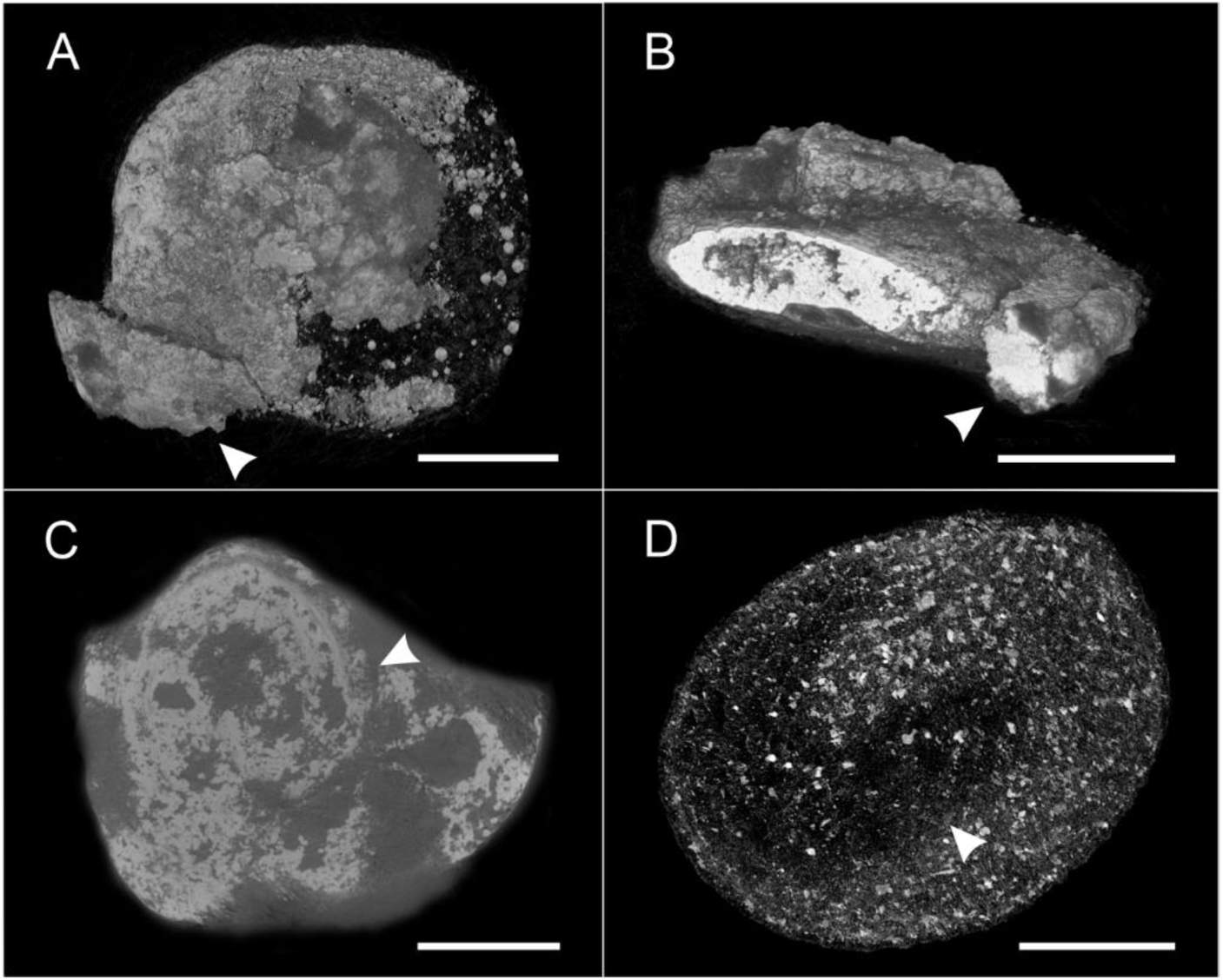
Micro-CT data of fossils showing diagnostic traits. **A-B.** *Hyoscyosperma daturoides* sp. nov., specimen BIN K453/133 (holotype); arrowheads point to the elaiosome attached to the seed tissue. **A.** Volume rendering, showing lateral view of the specimen. **B.** Digital transverse section to show the elaiosome attachment. **C.** Digital lateral section of broken specimen BIN K453/175; arrowhead indicates the clear gap where the coiled embryo is missing. **D.** *Solanum dorofeevii* sp. nov., digital lateral section of specimen MB.Pb1993/6802; note the gap where the curved embryo is missing (arrowhead). Bars: 0.5 mm.

### Morphospace of fossil and extant Solanaceae seeds

Our NMDS analysis showed that 12 out of the 14 morphological traits were significantly informative (P = 0.001–0.04), excluding the roundness and the perimeter/area ratio of the exotestal cells (Table S6). These characters were still retained because they are useful to separate clusters. A stress analysis (measuring goodness-of-fit) showed that two dimensions produce a stress value < 0.2, which indicates that the plot is interpretable. In our two-dimensional NMDS plot, the first axis (NMDS1) strongly separates taxa with and without compressed seeds (character 4), while the second axis (NMDS2) separates taxa mainly by the shape of exotestal anticlinal walls (character 1) and the position of the hilum (character 2; **Fig. 2**). The remaining characters contributed to the spread and clustering of the taxa, allowing us to identify distinct combinations of seed traits and areas in morphospace to serve as the foundation for classification of fossil seeds. In particular, we identified 10 clusters that included both living taxa and fossil specimens (circled in bold gray in **Fig. 2**; Table S1). We used these clusters to re-circumscribe fossil genera and species and describe new fossil genera and species (see below). Seven additional clusters (circled in thin gray in **Fig. 2**) included only fossils scored from the bibliography (i.e., no scored specimens); we elected not to include these in the taxonomic revision presented below.

**Figure 2.**
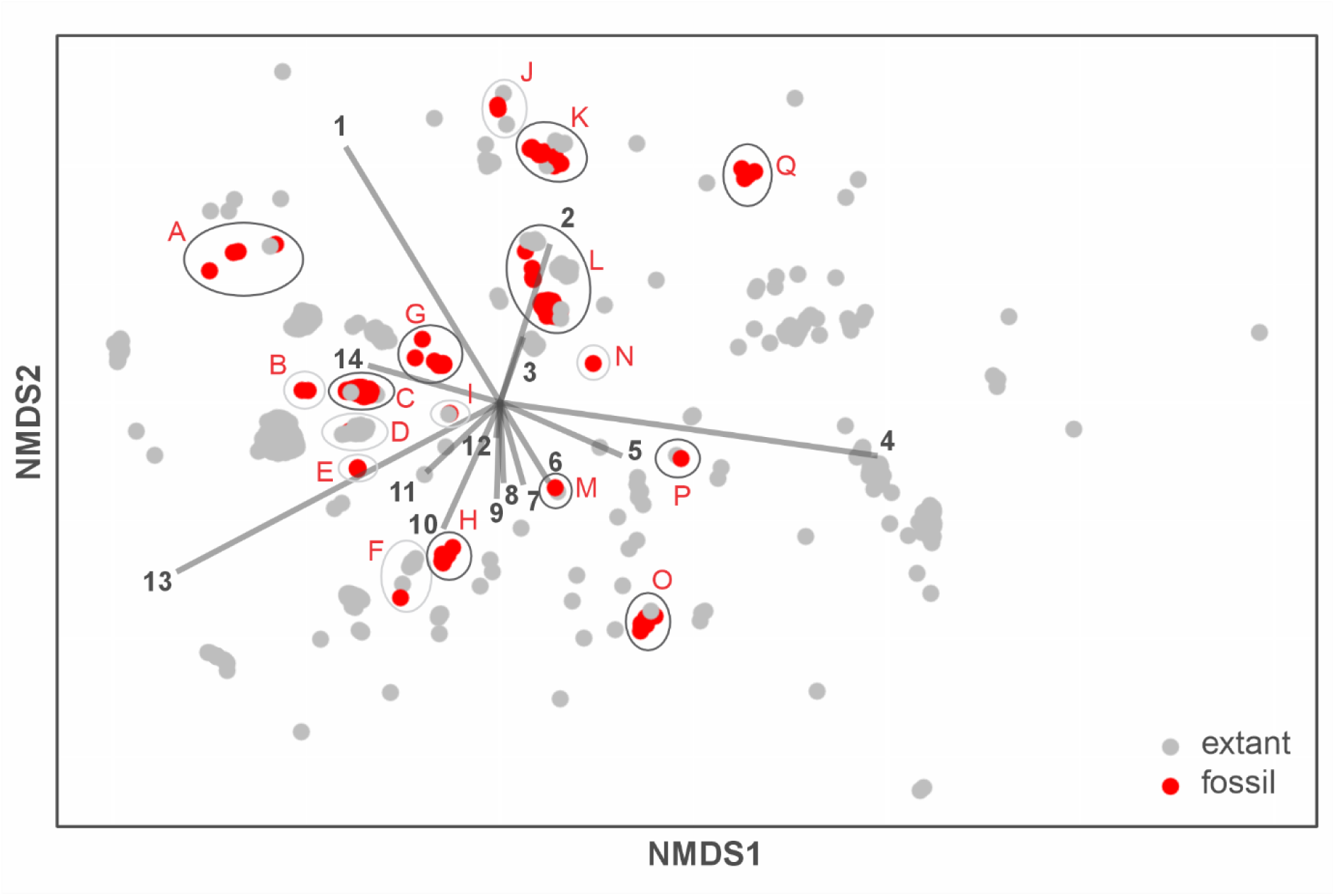
Non-metric multidimensional scaling (NMDS) analysis of 14 morphological characters from 110 fossil seeds and 354 extant species of Solanaceae, with 17 resulting groupings of fossil and extant taxa indicated. Dark grey circles denote clusters analyzed here to circumscribe new taxa or re-evaluate previously described genera and species. Light grey circles indicate putative taxa not treated further here because detailed analysis was not possible (characters scored only from patchy descriptions). Each character is indicated with a number, and the length of the line leading to the numbers gives the strength of the correlation with each NMDS axis. The characters are as follows: 1: shape of the anticlinal walls of the exotestal cells, 2: position of the hilum, 3: perimeter: area ratio of exotestal cells, 4: compression, 5: seed length/width ratio, 6: area of exotestal cells, 7: perimeter of exotestal cells, 8: seed length, 9: seed area, 10: embryo shape, 11: presence of wings, 12: roundness of exotestal cells, 13: presence of seed cavity, 14: presence of elaiosomes.

## TAXONOMIC DESCRIPTIONS

Order **Solanales** Bercht. & J. Presl, 1820 Family **Solanaceae** Juss., 1789, nom. cons. ***Hyoscyosperma*** Deanna & S.D.Sm., **gen. nov. –** Type: *Hyoscyosperma daturoides Generic diagnosis.* **–** Differing from any other known solanaceous seeds in the character combination of seeds compressed, usually with a hilum of lateral/sublateral position and a cavity at the hilar-chalazal end with a conspicuous elaiosome, and exotestal cell uniform, with anticlinal walls sinuate-cerebelloid on the entire seed surface.

*Etymology.* **–** Named after morphological similarity to the henbane seeds (*Hyoscyamus*; **Fig. 3D**). ***Hyoscyosperma daturoides*** Deanna & S.D.Sm., **sp. nov**.– Holotype: RUSSIA, Northern Caucasus, Rostov oblast, 47.762692, 42.746863, BIN K453/133 (Komarov Botanical Institute, St. Petersburg, Russia; **Fig. 3A**).

**Figure 3.**
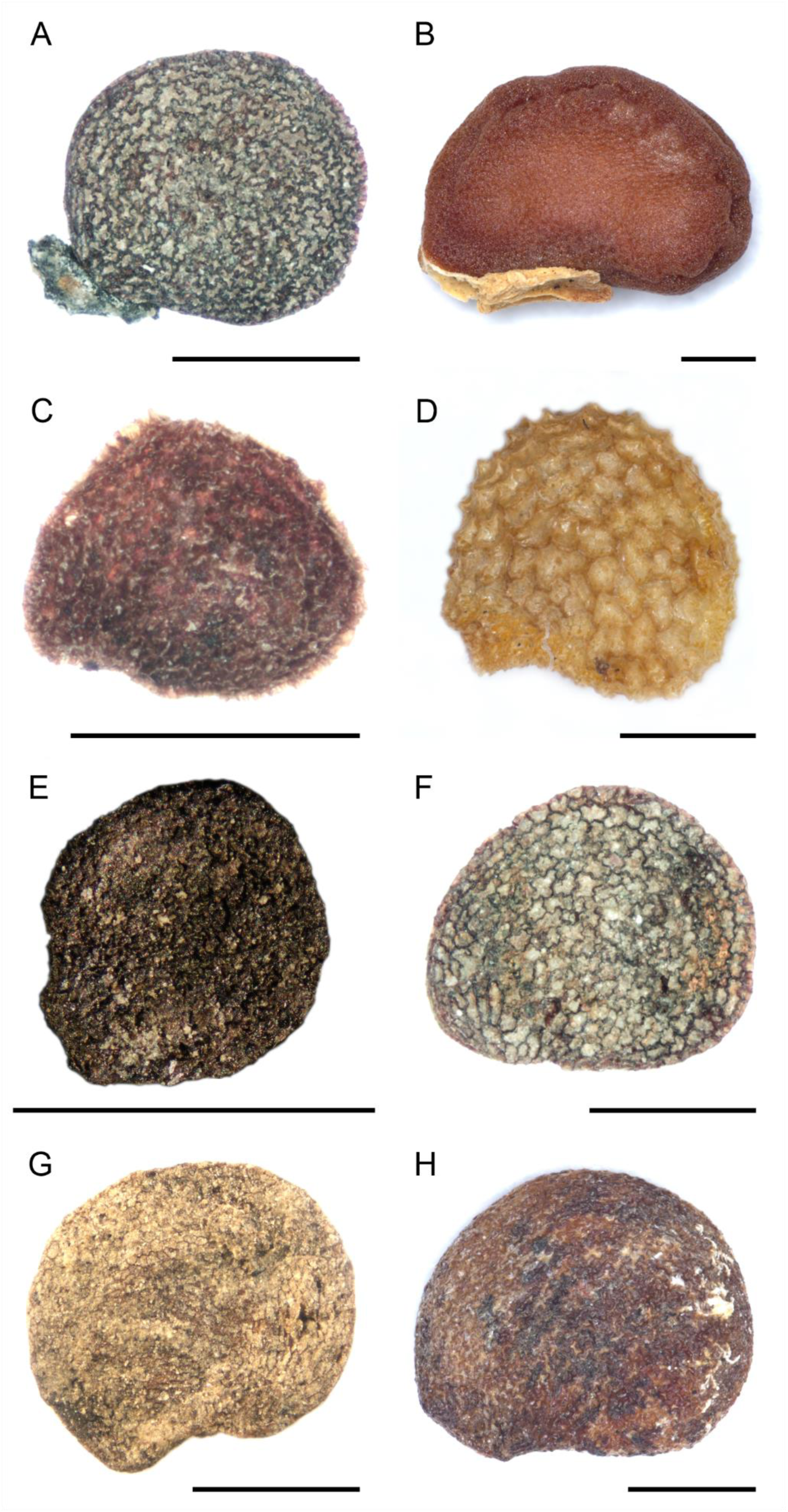
Fossil seeds of *Hyoscyosperma daturoides* Deanna & S.D.Sm. gen. et sp. nov.*, Solanum foveolatum* Negru and morphologically similar extant taxa. **A-D.** Specimens grouped in cluster A, *Hyoscyosperma daturoides*. **A.** Holotype of *H. daturoides*, BIN K453/133. **B.** Extant seed of *Datura metel* L., KIB7415. **C.** Paratype of *H. daturoides*, BIN H4044/71. **D. S**eed of the extant *Hyoscyamus niger* L. **E-F.** Specimens grouped in cluster C, *Solanum foveolatum*. **E.** Holotype of *S. foveolatum*, MB.Pb.1998/0434. **F.** Specimen BIN K587/31. **G.** Specimen MB.Pb.1993/0791. **H. S**eed of the extant *Anisodus luridus* Spreng. A, C, F photographs by A. V. Hvalj, B, D, E, G, H by R. Deanna. Bars: 1 mm.

*Species diagnosis*. – Same as for genus. *Specific description*. – Seeds compressed, 1.45 (1.19-1.71) mm long, 1.15 (0.97-1.54) mm wide, ratio length/width 1.28 (1.11-1.41). Hilum usually lateral/sublateral with a cavity at the hilar-chalazal end. Exotestal cells uniform in shape, area of 0.009 (0.004-0.014) mm^2^, perimeter of 0.55 (0.27-0.71) mm. Exotestal anticlinal walls sinuate-cerebelloid on the entire seed surface. The most important trait is the presence of an elaiosome (**Figs. 1A-B, 3A**) that relates this species to the genus *Datura* (**Fig. 3B**), although the combination of size, type of exotestal anticlinal walls, and measures of exotestal cells place *Hyoscyosperma daturoides* as most morphologically similar to the extant genus *Hyoscyamus* L. (**Fig. 3D**).

*Age and distribution.* – Late Oligocene to Quaternary, 27.8 – 0.01 Ma (including Dorofeev, 1957a). A complete stratigraphic range and a map showing localities are shown on **Fig. 4**.

**Figure 4.**
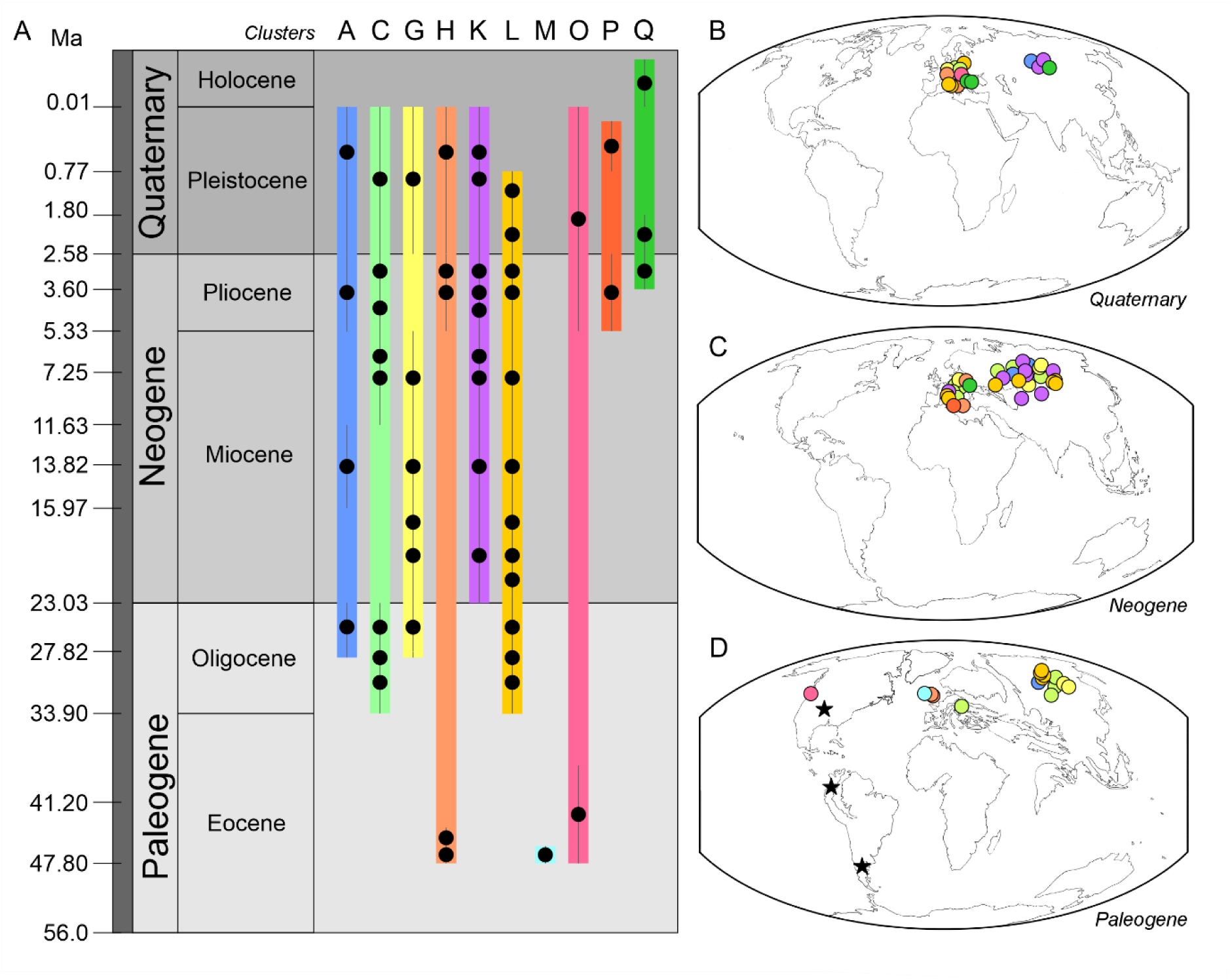
Temporal and geographical distribution of Solanaceae seed fossils analyzed according to the clusters identified. **A.** Temporal distribution (colored bars) and stratigraphic occurrences (black dots) of seed fossils of Solanaceae analyzed, plotted against the International Chronostratigraphic Chart (v. 2023/09). Letters on top represent the NMDS clusters: A containing *Hyoscyosperma daturoides;* C, *Solanum foveolatum*; G, *Solanum miocenicum*; H, *Solanispermum reniforme,* K, *Hyoscyamus undulatus*; L, *Solanoides dorofeevii*; M, *Albionites arnensis*; O, *Nephrosemen reticulatum*; P, *Capsicum pliocenicum*; Q, *Thanatosperma minutum*. **B-D.** Solanaceae seed occurrences, color-coded by clusters, are shown on paleogeographic reconstructions of the world corresponding to different geological periods. **B.** Quaternary. **C.** Neogene. **D.** Paleogene represented by the Eocene reconstruction of the world map. Stars represent occurrences of fruit fossils. Maps drawn by S. Monteschiesi from reconstructions by C. R. Scotese, PALEOMAP Project (Scotese & Dreher, 2012).

**Figure 5.**
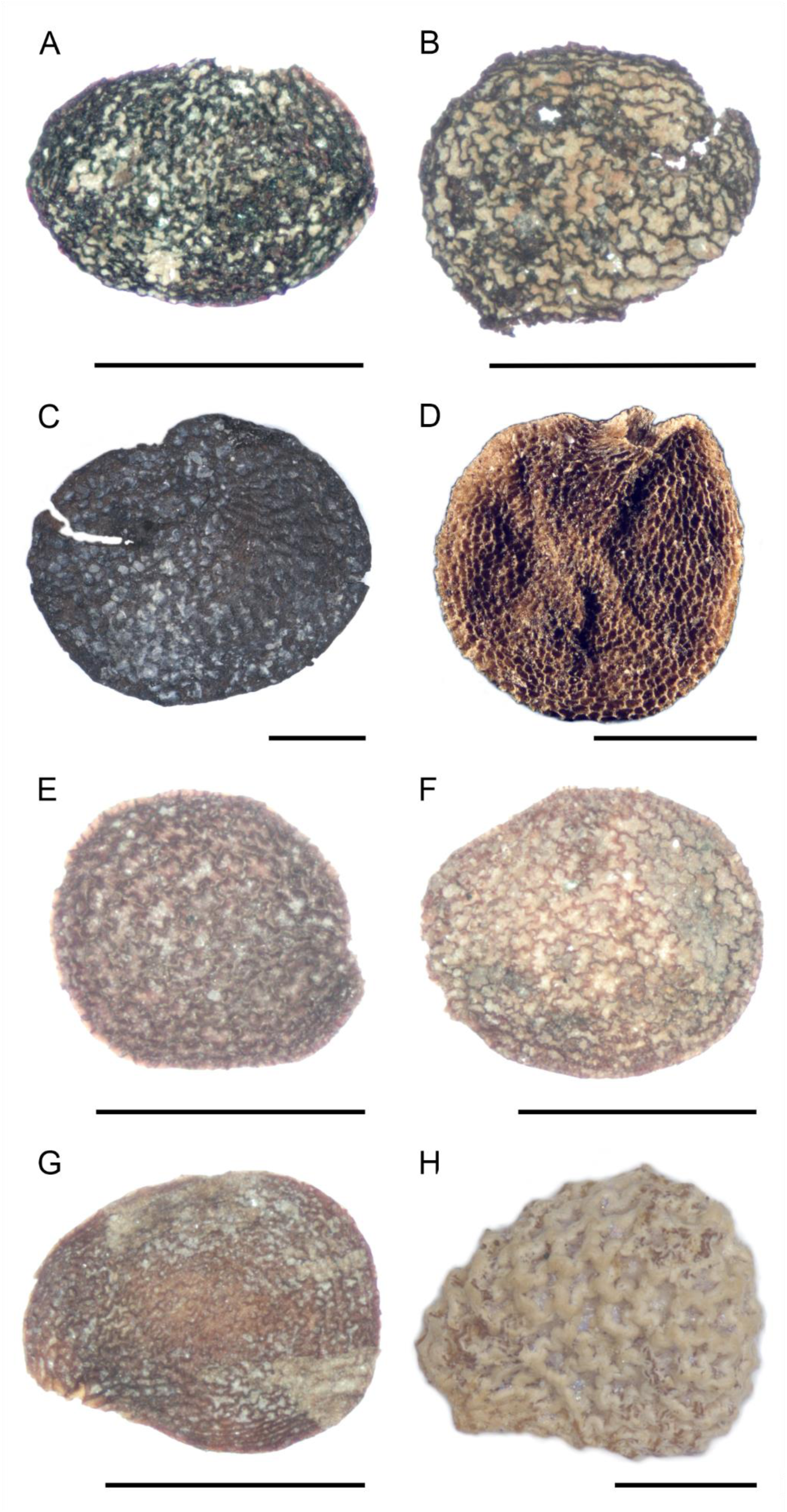
Fossils of *Solanum miocenicum* Deanna & S.D.Sm. sp. nov., *Solanispermum reniforme* M.Chandler*, Hyoscyamus undulatus* Deanna & S.D.Sm. sp nov., and morphologically similar extant taxa. **A-B.** Specimens grouped in cluster G, *Solanum miocenicum*. **A.** Holotype of *S. miocenicum*, BIN K528/50. **B.** Paratype of *S. miocenicum*, BIN H4177/82. **C-D.** Representative specimens of cluster H, *Solanispermum reniforme.* **C.** Paratype of *S. reniforme*, V-40891. **D.** New specimen included in *S. reniforme*, MB.Pb.1993/10027. **E-H.** Specimens grouped in cluster K, *Hyoscyamus undulatus*. **E.** Holotype of *H. undulatus* sp. nov., BIN H4895/49. **F.** Paratype of *H. undulatus*, BIN K587/30, previously identified as *Solanum dulcamara* L. (Dorofeev, 1977). **G.** Paratype of *H. undulatus,* BIN H4008/93. **H.** Seed of the extant *Hyoscyamus albus* L. A, B, E-G photographs by A. V. Hvalj, C, D, H by R. Deanna. Bars: 1 mm.

*Additional material analyzed.* – RUSSIA, West Siberia, Tomsk oblast, BIN H1594/44; KAZAKHSTAN, Soltüstık Qazaqstan oblysy, 54.30109, 68.346153, BIN H4044/71 (**Fig. 3C**); RUSSIA, West Siberia, Omsk oblast, 55.903982, 74.727913, BIN H554 (Komarov Botanical Institute, St. Petersburg, Russia).

*Etymology*. – Named after the resemblance to the Jimsonweed genus (*Datura*) given the presence of an elaiosome of the same size, shape, and attachment (**Figs. 1A-B**, **3A-B**).

*Remarks.* – This species comprises all fossil seed taxa grouped in cluster A from the NMDS analysis (**Fig. 2**). This cluster includes four fossils and two extant seeds with a particular combination of traits (Table S1). Although *Datura* L. (**Fig. 3B**) is the only genus in the Solanaceae with seeds containing an elaiosome of the same shape, size, and attachment as the holotype of *Hyoscyosperma daturoides* (BIN K453/133, **Fig. 3A**), *Datura* seeds are much larger, and the exotestal cell characteristics are distinct from this entire cluster of fossils. These traits scored on the fossils are consistent with the extant genus *Hyoscyamus* (**Fig. 3D**), especially the uniform shape and size of the exotestal cells on the entire seed surface and the overall size and shape of the seed.

The holotype of *Hyoscyosperma daturoides* (BIN K453/133) was previously identified as *Physalis pliocenica* Szafer by Dorofeev (1957a) based on the seed’s shape and testa. However, this seed differs from the species description (Szafer, 1946) in its actual shape and the presence of an elaiosome. *Physalis pliocenica* was initially described by Szafer (1946) including five seeds and a damaged fruiting calyx of c. 15 mm diameter but Szafer (1946) did not propose systematic affinities, and we found one of the two seeds imaged by Szafer (1946, pl. 13, fig. 23) clustered with the fossil 14183-28 (van der Burgh, 1987) and two extant taxa (Cluster J; **Fig. 2**).

***Solanum foveolatum*** Negru, Meot. Fl. Sev.-Zap. Prichernomor. 128. 1986, ***emend.*** – Holotype: GERMANY, Bauersberg, MB.Pb.1998/0434 (Museum für Naturkunde Berlin, Germany; Negru 1986, pl. 18, figs. 2-7; **Fig. 3E**).

*Original species diagnosis.* – “Samen im Umriss fast kreisförmig, an der Basis ein wenig grubig eingetieft und abgestutzt, fast eben über die Seiten. Testa relativ dünnwandig, von auβen grubig. Grübchen rundlich-vieleckig, tief, mit ungleich hohen, dünnen, gleichförmigen Wänden. Am Hilum die Grübchen viel kleiner, eng [Seeds are almost circular in outline, slightly pitted and truncated at the base, almost flat across the sides. Testa relatively thin-walled, pitted on the outside. Pits are roundish-polygonal, deep, with unevenly high, thin, uniform walls. At the hilum, the pits are much smaller and narrow.]” (Negru 1986, p. 128-129).

*Emended species diagnosis.* – Differing from any other known solanaceous seeds in the character combination of seeds compressed, almost circular in outline, always with lateral/sublateral hilum, and with a cavity at the hilar-chalazal end, and exotestal cells uniform in shape and size on the entire seed surface, with sinuate-cerebelloid and thin anticlinal walls, except towards the hilum where the cells are smaller.

*Species description.* – Seeds compressed, 1.56 (0.83-2.34) mm long, 1.35 (0.804-2.02) mm wide, almost circular in outline, length/width ratio of 1.15 (1-1.37). Hilum lateral/sublateral with a cavity at the hilar-chalazal end (**Fig. 3E-G**). Without elaiosomes or wings. Exotestal cells uniform in shape and size on the entire seed surface, except towards the hilum where the cells are smaller, exotestal cells area of 0.0106 (0.002-0.020) mm^2^, perimeter of 0.55 (0.24-1.08) mm at the center of the seed surface. Exotestal anticlinal walls sinuate-cerebelloid and thin. Embryo shape unknown.

*Age and distribution.* – Early Oligocene to Pleistocene, 33.9 – 0.01 Ma (including Negru 1986; Dorofeev 1960, 1977; Mai & al., 1963; Mai, 1997; Basilici & al., 1997; Mai, 2000, 2010). A complete stratigraphic range and geographic distribution are shown in **Fig. 4**.

*Additional material analyzed.* – RUSSIA, Ural, Bashkiria, 54.834631, 56.397617, BIN K390; RUSSIA, Ural, Bashkiria, 54.753123, 54.883264, BIN K587/31 (**Fig. 3F**); RUSSIA, West Siberia, Tomsk oblast, 58.328408, 79.403543, BIN H4759; BELARUS, Kholmyech, 52.158587, 30.623945, BIN K690; RUSSIA, West Siberia, Tomsk oblast, 60.100319, 83.209543, BIN K536; RUSSIA, West Siberia, Novosibirsk oblast, 54.952243, 82.814527, BIN H4308; RUSSIA, West Siberia, Novosibirsk oblast, 55.266820, 79.146436, BIN H1818; RUSSIA, West Siberia, Tomsk oblast, 60.108504, 83.242754, BIN H4152; RUSSIA, West Siberia, Tomsk oblast, 57.021166, 82.071869, BIN H1328; RUSSIA, West Siberia, Tomsk oblast, 56.726710, 86.453605, BIN H1120; RUSSIA, West Siberia, Omsk oblast, 56.648865, 73.015980, BIN H3564 (Komarov Botanical Institute, St. Petersburg, Russia); GERMANY, Hartau b. Zittau, MB.Pb.1993/7605; GERMANY, Mahlis, MB.Pb. 1993/9655B; GERMANY, Nordhausen, MB.Pb.1993/6831; GERMANY, Berga b. Sangerhausen, MB.Pb.1993/5546; GERMANY, Horka, MB.Pb.1993/2220, GERMANY, GERMANY, Rippersroda b. Arnstadt, MB.Pb.1993/0791 (**Fig. 3G**); Braunsbedra b. Merseburg, MB.Pb.1993/10814 (Museum für Naturkunde Berlin, Germany); ITALY, Sento-S7a, MRSN-P/345-CCN4332 (CENOFITA collection, managed by the Regional Museum of Natural Sciences of Turin, Turin, Italy).

*Remarks.* – This species comprises all fossil seed specimens grouped in cluster C of the NMDS analysis (**Fig. 2**), which groups 20 fossil seeds and two extant taxa, the genus *Anisodus* Spreng. (**Fig. 3H**) from the Hyoscyameae clade and the genus *Cuatresia* Hunz. from the Physalideae clade (Table S1). Both genera are easily recognized as members of the berry clade within subfamily Solanoideae by the flattened seeds always with a lateral/sublateral hilum and a cavity at the hilar-chalazal end, as with all the fossils included in the cluster. These morphological traits support the placement of this fossil species in the berry clade, but there is no further evidence found to assign it to a particular extant genus. However, we cannot exclude the possibility that it belongs to *Solanum* L. and so, in order to keep a stable nomenclature, we have decided to retain the name already in use (Gumbel & Mai, 2002), *Solanum foveolatum*.

Most seed fossils of the nightshade family have been identified as extant taxa of the genera *Solanum* or *Physalis*. *Solanum foveolatum* is no exception, we found two fossils previously identified as *Alkekengi officinarum* Moench (= *Physalis alkekengi* L., BIN K390 and BIN K587/31, **Fig. 3F**; Dorofeev 1960, 1977), one fossil as *Physalis plioceniza* Szafer (MB.Pb.1993/7605; Mai, 2000), and three fossils as *Solanum dulcamara* L. (MB.Pb.1993/0791, MB.Pb.1993/10814, and MRSN-P/345-CCN4332; **Fig. 3G**; Mai & al., 1963; Mai, 2010; Basilici & al., 1997) clustered with the type specimen of *S. foveolatum* based on heterogeneous shape of the exotestal cells from the center to the margins of the seed, perimeter, roundness, and area of the exotestal cells. Two other fossils (MB.Pb.1993/2220 and MB.Pb.1993/7605; Mai 1997, 2000) were previously reported as *Physalis pliocenica,* most likely due to the characteristic seed traits of the berry clade.

***Solanum miocenicum*** Deanna & S.D.Sm., **sp. nov.** – Holotype: RUSSIA, West Siberia, Omsk oblast, 57.734521, 71.173771, BIN K528/50 (Komarov Botanical Institute, St. Petersburg, Russia; **Fig. 5A**).

*Species diagnosis.* – Differing from any other known solanaceous seeds in the character combination of seeds compressed, round to slightly elongated, hilum lateral/sublateral, without a cavity at the hilar-chalazal end, and exotestal cells uniform in shape, with anticlinal walls strongly cerebelloid on the entire seed surface.

*Species description.* – Seeds compressed, round, slightly elongated, 1.46 (1.25-1.87) mm long, 1.16 (0.93-1.41) mm wide, ratio length/width 1.26 (1.03-1.41). Hilum lateral/sublateral, without a cavity at the hilar-chalazal end. Exotestal cells uniform in shape, area of 0.012 (0.006-0.016) mm^2^, perimeter of 0.68 (0.43-0.79) mm. Exotestal anticlinal walls strongly cerebelloid on the entire seed surface (**Fig. 5A-B**). Embryo shape unknown.

*Age and distribution.* – Late Oligocene to Pleistocene, 27.82 – 0.01 Ma (including Dorofeev, 1963). A complete stratigraphic range and a map showing localities are shown on **Fig. 4**.

*Additional material analyzed.* – RUSSIA, West Siberia, Omsk oblast, 56.671976, 74.706109, BIN H4177/82 (**Fig. 5B**); RUSSIA, West Siberia, Novosibirsk oblast, 53.839196, 77.443347, BIN H2373; RUSSIA, 61.44233, 83.864155, West Siberia, Omsk oblast, 57.693604, 71.393553, BIN K527; West Siberia, RUSSIA, West Siberia, Khanty-Mansi Autonomous Okrug–Yugra, BIN H1344; RUSSIA, West Siberia, Novosibirsk oblast, 56.049558, 84.198969, BIN H2000 (Komarov Botanical Institute, St. Petersburg, Russia); GERMANY, Mahlis, MB.Pb.1993/9655; GERMANY, Bröthen b. Hoyerswerda, MB.Pb.1993/4558 (Museum für Naturkunde Berlin, Germany).

*Etymology.* – The name of this species relates to its stratigraphic distribution, with most of its fossil specimens coming from the Miocene.

*Remarks.* – This species comprises all fossil seed taxa clustering in cluster G of the NMDS analysis (**Fig. 2**), which includes eight seed fossils (Tables S1, S3). The combination of their morphological traits (Table S3) supports the placement of this fossil species in the berry clade, within *Solanum,* because of the absence of a cavity at the hilar-chalazal end, shared with several *Solanum* extant species. *Solanum miocenicum* includes two previously published fossils assigned to the genus *Solanum* (K528/50, K527, Dorofeev, 1963) because of the morphological similarities with the extant genus.

***Solanispermum reniforme*** M.Chandler, Bull. Brit. Mus. (Nat. Hist.), Geol. 3(3): 118. 1957, ***emend.*** – Holotype: UNITED KINGDOM, Bovey Tracey, GSM76684 (=GSM1805; British Geological Survey, Keyworth, Nottinghamshire, United Kingdom; Chandler 1957, pl. 17, figs. 189-191; Reid & Reid 1910, pl. 16, fig. 72).

*Original species diagnosis.* “Seeds are transversely oval or reniform in outline, occasionally hooked, and surface normally with coarse, interrupted, sinuous rugosities or tubercles which produce a pitted effect in places. An outer coat, rarely preserved, shows ‘pits’ with sinuous outlines. The rugose coat shows fine striae at right angles to the tubercles. Splitting along the striae on drying produces a fibrous effect. Inner coat springy, formed of equiaxial cells. Maximum diameter of seeds about 3.5-4.8 mm” (Chandler, 1957).

*Emended species diagnosis.* – Differing from any other known solanaceous seeds in the character combination of seeds compressed, oval to reniform in outline, always with lateral/sublateral hilum and with a cavity at the hilar-chalazal end, and exotestal cells varying in shape, irregularly circular in the center of the seed surface, rectangular towards the margins and the hilum, with straight to slightly sinuate anticlinal walls showing pits. It also differs by the seed coat rugose, striate, the striae crossing the rugosities at right angles.

*Species description.* – Seeds compressed, 2.35 (1.54-3.65) mm long, 1.98 (1.45-3.04) mm wide, oval to reniform in outline, length/width ratio of 1.17 (1.02-1.25). Hilum always lateral/sublateral, with a cavity at the hilar-chalazal end (**Fig. 5C-D**). Without elaiosomes or wings. Exotestal cells non-uniform in shape, irregularly circular in the center of the seed surface, rectangular towards the margins and the hilum, with straight to slightly sinuate anticlinal walls showing pits of 0.05-0.1 mm diam. Exotestal cells area of 0.005 (0.002-0.012) mm^2^, perimeter of 0.33 (0.19-0.56) mm at the center of the seed surface. Seed coat rugose, 0.1 mm thick, striate, the striae crossing the rugosities at right angles. Splitting along the striae on drying produces a fibrous effect. These ‘fibres’ are formed of fine equiaxial cells of 0.012 mm diam, occurring as several layers in the coat. Spongy coat within, 0.4 mm thick, composed of dense soft parenchyma. Embryo shape unknown.

*Age and distribution.* – Late Ypresian/early Lutetian to Quaternary, 48-0.01 Ma (including Reid & Reid, 1910; Chandler, 1957; Mai & Walther, 1988; Martinetto, 1995). A complete stratigraphic range and a map showing localities are shown in **Fig. 4**.

*Additional material analyzed.* – UNITED KINGDOM, Arne, Poole Formation Sandbanks, Poole Formation, NHMUK V-40891 (**Fig. 5C**); UNITED KINGDOM, Bovey Tracey, GSM76684 (=GSM1805; British Geological Survey, Keyworth, Nottinghamshire, United Kingdom); UNITED KINGDOM, Branksome Dene, Bournemouth Freshwater Beds, Branksome Sand Formation, NHMUK V-42019 (Natural History Museum, London, United Kingdom); ITALY, Bucine, MRSN-P/345-CCN6065a; ITALY, Ca’ Viettone, MRSN-P/345-CCN4328a; ITALY, Ca’ Viettone, MRSN-P/345-CCN4328b (CENOFITA collection, managed by the Regional Museum of Natural Sciences of Turin, Turin, Italy); GERMANY, Berga b. Sangerhausen, MB.Pb.1993/5544; GERMANY, Berga b. Sangerhausen, MB.Pb.1993/5544a; GERMANY, Seeberg, MB.Pb.1993/10027 (**Fig. 5D**; Museum für Naturkunde Berlin, Germany).

*Remarks.* – This species comprises all fossil seed taxa grouped in cluster H resulting from the NMDS analysis (**Fig. 2**), which included only nine fossils. The distinctness of this taxon from any extant supports its recognition as a fossil genus, as it was initially described by Chandler (1957), and later confirmed by Chandler (1962, 1963). This fossil species now also includes occurrences from Germany and Italy, which better represent the variability of the morphological traits and enlarge the stratigraphical range of this taxon towards the present. The combination of the flattened campylotropous seeds with curved outline and marginal hilar cavity strongly suggests that this fossil taxon belongs to Solanaceae, and most likely to the Solanoideae clade. However, the combination of a seed coat with multiple layers and mostly straight exotestal anticlinal walls does not match with any extant genera in the family, and so, this fossil taxon is kept as an extinct genus. *Solanispermum reniforme* is the earliest seed in the fossil record of the family (Särkinen & al., 2018) and extends from the Eocene through the Holocene (**Fig. 4)**.

***Hyoscyamus undulatus*** Deanna & S.D.Sm., **sp. nov.** – Holotype: RUSSIA, West Siberia, Novosibirsk oblast, 55.814554, 84.524896, BIN H4895/49 (Komarov Botanical Institute, St. Petersburg, Russia; **Fig. 5E**).

*Species diagnosis.* – Differing from any other known solanaceous seeds in the character combination of seeds compressed, round to oblong, hilum terminal, slightly protruding, without a cavity at the hilar-chalazal end, and exotestal cells uniform in shape, with anticlinal walls sinuate-cerebelloid on the entire seed surface.

*Species description.* – Seeds compressed, round to oblong, 1.59 (1.06-2.05) mm long, 1.26 (1.03-1.59) mm wide, ratio length/width 1.26 (1.03-1.59). Hilum terminal, slightly protruding (**Fig. 5G**), without a cavity at the hilar-chalazal end. Exotestal cells uniform in shape, area of 0.01 (0.003-0.016) mm^2^, perimeter of 0.51 (0.30-0.67) mm. Anticlinal walls of exotestal cells sinuate-cerebelloid on the entire seed surface (**Fig. 5E-G**). Embryo shape unknown.

*Age and distribution.* – Early Miocene to Quaternary, 23.03-0.01 Ma (including Dorofeev, 1956, 1957a, 1957b, 1963, 1966, 1977). A complete stratigraphic range and a map showing localities are shown on **Fig. 4**.

*Additional material analyzed.* – RUSSIA, Ural, Bashkiria, 54.753123, 54.883264, BIN K587/30 (**Fig. 5F**); RUSSIA, Northern Caucasus, Rostov oblast, 47.762692, 42.746863, BIN K453; RUSSIA, Volga region, Volgograd oblast, 48.484398, 44.786488, BIN K345; RUSSIA, West Siberia, Tomsk oblast, 56.030961, 83.895555, BIN H135; RUSSIA, West Siberia, Altay kray, 53.354407, 82.237467, BIN H1280; GEORGIA, Abkhazia, Duab, 42.828644, 41.504082, BIN K531; RUSSIA, West Siberia, Tomsk oblast, 56.343530, 84.087431, BIN K388; KAZAKHASTAN, Pavlodar oblysy, 53.997642, 76.373463, BIN H4008/93 (**Fig. 5G)**; RUSSIA, Volga region, Tataria, 55.448199, 51.40682, BIN K366; RUSSIA, West Siberia, Omsk oblast, 57.495903, 73.844962, BIN H4390 (Komarov Botanical Institute, St. Petersburg, Russia); POLAND, Kroscienko, MB.Pb.1993/7754; GERMANY, Nordhausen, MB.Pb.1993/6832 (Museum für Naturkunde Berlin, Germany).

*Etymology.* – The specific epithet *undulatus* refers to the sinuate-cerebelloid anticlinal walls of the exotestal cells of the entire surface of the seeds.

*Remarks.* – This species comprises all fossil seed taxa clustering in cluster K result of the NMDS analysis, which includes 13 fossils grouped with four extant species of the genus *Hyoscyamus* (**Fig. 5H**) and one species of *Lycianthes* Hassl. Fossil specimens once identified as *Hyoscyamus niger* L. (BIN K453; Dorofeev, 1957a, 1966), *Solanum dulcamara* L. (K587/30; Dorofeev, 1977), *Solanum nigrum* L. (BIN K345, BIN K366; Dorofeev, 1956, 1957b), and *Physalis* (BIN K388; Dorofeev, 1963), are now reassigned here based on a unique combination of traits. The terminal, protruding hilum and the distinctly sinuate-cerebelloid exotestal anticlinal walls are diagnostic for this species and match the defining characteristics of *Hyoscyamus* (**Fig. 5H**; Kaya & al., 2016).

Among the defining morphological features, the terminal hilum is particularly conspicuous. This character is otherwise only shared with *Thanatosperma minutum* and *Hyosycosperma daturoides* within the broader fossil dataset. However, it clearly differs from *Hyosycosperma* by lacking an elaiosome and from *Thanatosperma* by its sinuate-cerebelloid exotestal anticlinal walls, in contrast to the straight walls of the latter. These differences confirm that this is a distinct species, morphologically alike and with a similar distribution to the extant *Hyoscyamus*. This taxon is among the most widely distributed seed species in the Solanaceae fossil record, after *Solanoides dorofeevii* and *Nephrosemen reticulatum*. Its occurrences span a broad geographic area, from West Siberia and Kazakhstan to Germany (**Fig. 4**).

***Solanoides*** Deanna & S.D.Sm., **gen. nov.** – Type: *Solanoides dorofeevii Generic diagnosis.* **–** Differing from any other known solanaceous seeds in the character combination of seeds compressed, hilum lateral/sublateral, without a cavity at the hilar-chalazal end, embryo curved, and exotestal cells varying in shape, irregular to round in the center of the seed surface, rectangular towards the margins, with anticlinal walls sinuate to slightly cerebelloid on the entire seed surface.

*Etymology.* **–** Named for its similar morphology to many other genera across the breadth of diversity in Solanaceae.

***Solanoides dorofeevii*** Deanna & S.D.Sm., **sp. nov.** – Holotype: RUSSIA, Northern Caucasus, Krasnodar kray, 44.515202, 39.730509, BIN K530/21 (Komarov Botanical Institute, St. Petersburg, Russia; Dorofeev, 1964; **Fig. 6A**).

**Figure 6.**
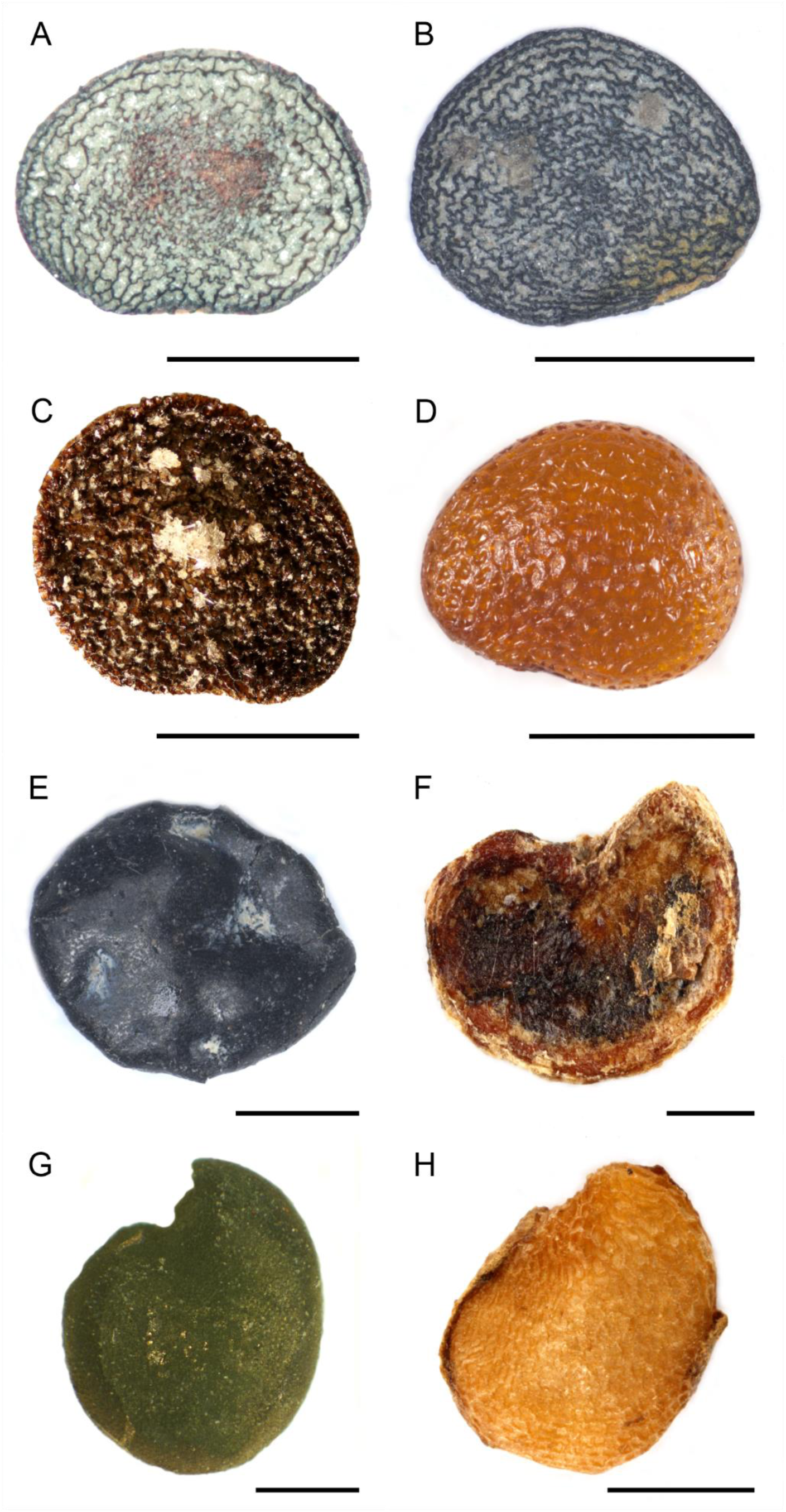
Fossils of *Solanoides dorofeevii* Deanna & S.D.Sm. gen. et sp. nov.*, Albionites arnensis* Deanna & S.Knapp. gen. et comb. nov.*, Nephrosemen reticulatum* Manchester, and morphologically similar extant taxa. **A-D.** Specimens grouped in cluster L, *Solanoides dorofeevii*. **A.** Holotype of *S. dorofeevii*, BIN K530/21. **B.** Paratype of *S. dorofeevii*, MRSN-P/345-CCN4324. **C.** Paratype of *S. dorofeevii,* MB.Pb.1993/5546B**. D. S**eed of the extant *Calliphysalis carpenteri* (Riddell) Whitson. **E-F.** Representative specimens of cluster M, *Albionites arnensis.* **E.** Holotype of *A. arnensis*, V-40898 (Chandler, 1962). **F.** Seed of the extant *Lycianthes oliveriana* (Lauterb. & Schum.) Bitter. **G-H.** Specimens grouped in cluster O, *Nephrosemen reticulatum.* **G.** Holotype of *N. reticulatum*, UF6500. **H.** Seed of the extant *Trozelia umbellata* Raf. (A photograph by A. V. Hvalj, B-H by R. Deanna). Bars: 1 mm.

*Species diagnosis.* – Same as for genus.

*Specific description*. – Seeds compressed, round to oblong-elongated, 1.51 (0.88-2.32) mm long, 1.22 (0.65-1.61) mm wide, ratio length/width 1.24 (1-1.7). Hilum lateral/sublateral, without a cavity at the hilar-chalazal end. Exotestal cell varying in shape, irregular to round in the center of the seed surface, rectangular towards the margins. Exotestal anticlinal walls sinuate to slightly cerebelloid on the entire seed surface. Exotestal cells area of 0.012 (0.003-0.025) mm^2^, perimeter of 0.58 (0.23-1.25) mm at the center of the seed surface. (**Fig. 6A-C**). Embryo curved (**Fig. 1D**).

*Age and distribution.* – Early Oligocene to early Pleistocene, 33.9 – 0.77 Ma (including Reid & Reid, 1910; Reid, 1923; Dorofeev, 1951, 1955, 1964, 1988; Szafer, 1961; Mai & Walther, 1988; Carvallo & Martinetto, 2001; Kovar-Eder & al., 2006; Martinetto, & Festa, 2013; Niccolini & al. 2022). A complete stratigraphic range and a map showing localities are shown on **Fig. 4**.

*Additional material analyzed.* – KAZAKHSTAN, Pavlodar, 53.997642, 76.373463, H4008; RUSSIA, West Siberia, Omsk oblast, 57.131842, 74.603660, H4220; RUSSIA, Novosibirsk oblast, 54.710437, 77.821560, H4736; RUSSIA, West Siberia, Tomsk oblast, 60.108504, 83.242754, H4148; RUSSIA, Central Black Earth region, Tambov oblast, 51.91591, 42.469724, K266; RUSSIA, West Siberia, Novosisibirsk oblast, 56.241528, 77.358989, H2727; RUSSIA, West Siberia, Novosibirsk oblast, 54.020703, 77.512922, H4000; RUSSIA, West Siberia, Yamalo-Nenets autonomous okrug, 64.836188, 82.933646, H2993; RUSSIA, West Siberia, Tomsk oblast, 60.100319, 83.209543, K518; RUSSIA; West Siberia, Tyumen oblast, Khanty-Mansy autonomous district, 61.425672, 82.885153, H1341; UKRAINE, Odesa oblast, Odesa, 46.410147, 30.758181, K432 (Komarov Botanical Institute, St. Petersburg, Russia); POLAND, Kroscienko, MB.Pb.1993/7753; GERMANY, Berzdorf, MB.Pb.1993/8849; GERMANY; Berga b. Sangerhausen, MB.Pb.1993/5545; GERMANY; Berga b. Sangerhausen, MB.Pb.1993/5546B **(Fig. 6C);** THE NETHERLANDS, Öbel, MB.Pb.1993/11406; THE NETHERLANDS, Rusell-Tiglia Egypte / Tegelen, MB.Pb.1993/6802 (**Fig. 1D**; Museum für Naturkunde Berlin, Germany); THE NETHERLANDS, Limburg, Tagelen, RGM793435 (Naturalis Biodiversity Center, Leiden, The Netherlands); ITALY, Buronzo, MRSN-P/345-CCN4334; ITALY, Castelletto Cervo II, MRSN-P/345-CCN3059a; ITALY, Castelletto Cervo II, MRSN-P/345-CCN1966B; ITALY, Castelletto Cervo II, MRSN-P/345-CCN3059b; ITALY, Ciabot Cagna, Corneliano d’Alba, MRSN-P/345-CCN4324 **(Fig. 6B)**; ITALY, Arda, MRSN-P/345-CCN4321 (CENOFITA collection, managed by the Regional Museum of Natural Sciences of Turin, Turin, Italy).

*Etymology.* – Named in honour of P.I. Dorofeev, Russian paleontologist, prolific collector of seed fossils, and author of many scientific publications, including on seed fossils of Solanaceae (e.g., Dorofeev, 1951, 1955, 1957a, 1957b, 1963, 1964, 1966, 1977).

*Remarks.* – This species comprises all fossil seed taxa grouped in cluster L from the NMDS analysis (**Fig. 2**), which includes 27 seed fossils and 24 extant taxa belonging to 16 genera across Solanaceae. This taxon encompasses fossils previously assigned to a wide array of genera within the Solanoideae clade, including *Physalis*, *Alkekengi* Moench, *Solanum*, *Hyoscyamus*, and *Scopolia* (Reid & Reid, 1910; Reid, 1923; Dorofeev, 1951, 1955, 1964, 1988; Szafer, 1961; Mai & Walther, 1988; Carvallo & Martinetto, 2001; Kovar-Eder & al., 2006; Martinetto, & Festa, 2013; Niccolini & al. 2022). Interestingly, the extant species grouped with this fossil taxon also span disparate genera—including *Nolana* L.f., *Benthamiella* Speg., and *Vestia* Willd.—that are not part of the Solanoideae clade. This suggests a potential case of morphological convergence or retention of plesiomorphic traits within seed coat architecture. The wide range of extant genera to which these fossils were previously assigned, in addition to the broad taxonomic span of the extant taxa in our cluster analysis, highlights the morphological variation of this fossil species.

*Solanoides dorofeevii* also stands out as the most geographically widespread seed species in the Solanaceae fossil record. Its known distribution ranges from Kazakhstan, and West Siberia to France. Although variable in terms of size and shape, this species is characterized by a unique combination of morphological traits: a hilum in lateral to sublateral position, the lack of a cavity at the hilar-chalazal end, the exotesta composed of variously-shaped cells with sinuate to slightly cerebelloid anticlinal walls, and a curved embryo. This suite of features distinguishes *S. dorofeevii* from all other fossil taxa.

***Albionites*** Deanna & S.Knapp, **gen. nov.** – Type: *Albionites arnensis* (=*Solanum arnense*)

*Generic diagnosis.* **–** Differing from any other known solanaceous seeds in the character combination of seeds globose, slightly laterally compressed towards the micropylar end, hilum lateral/sublateral, with a small cavity at the hilar-chalazal end, and exotestal cells uniform in shape, with anticlinal walls cerebelloid, with five to seven digitations.

*Etymology.* **–** Named after Albion, the literary name for the south of England, where this fossil was found.

***Albionites arnensis*** (M.Chandler) Deanna & S.Knapp, **comb nov.** ≡ *Solanum arnense* M.Chandler, Lower Tert. Fl. S. England 2: 141. 1962. – Holotype: UNITED KINGDOM, Arne, Poole Formation, NHMUK V40898 (Natural History Museum, London, United Kingdom; Chandler, 1962, pl. 22, figs. 12-13, text-fig. 22; **Fig. 6E**).

*Original species diagnosis.* – Hilum gaping not deeply rimmed, surface smooth, formed of conspicuously digitate cells, about 0-085 in diameter, with raised double outlines. Digitations about five or seven to one cell. Maximum diameter, 2-5 to 3 mm (Chandler, 1962).

*Species description.* – Seeds globose, obliquely oboval, slightly laterally compressed towards the micropylar end, bisymmetric about a plane passing through hilum and micropyle, curved, 2.65 mm long, 2.35 mm wide, length/width ratio of 1.13. Hilum large, lateral/sublateral, with a small cavity at the hilar-chalazal end separated from the main seed-cavity by a small, curved partition (**Fig. 6E**). Seed surface smooth, shining, black (as preserved). Exotestal cells uniform in shape, area of 0.003 mm^2^, perimeter of 0.18 mm. The outlines of these cells are double and often slightly raised. Exotestal anticlinal walls cerebelloid, with five to seven finger-like projections. These projections may be long and are sometimes curved or slightly bifid. In section, the exotesta is formed of fluted columns with the flutings corresponding with the digitations of the cells. An underlying testa layer is present, formed of angular cells in surface view.

*Age and distribution.* – Eocene, late Ypresian / early Lutetian, 48-46 Ma (including Chandler, 1962).

A complete stratigraphic range and a map showing localities are shown on **Fig. 4**.

*Remarks.* – This species comprises all fossil seed taxa grouped in cluster M of the NMDS analysis (**Fig. 2**), including a single fossil and the extant Australasian *Lycianthes oliveriana* (Lauterb. & Schum.) Bitter (**Fig. 6E-F**). As Chandler (1962) discussed, this fossil agrees with Solanaceae in the curved form, lateral hilar cavity, and cerebelloid exotestal cells with raised anticlinal walls. Her placement of the fossil in *Solanum* by comparison to the extant species *Solanum marginatum* L.f. (Chandler, 1962), however, cannot be upheld, as the morphology of the distinctly flattened seeds of that species do not correspond to this fossil. This extinct genus is different from all remaining fossils examined by the combination of the lack of general compression, the presence of a hilar cavity, and the cerebelloid exotestal anticlinal walls.

***Nephrosemen reticulatum*** Manchester, Paleontogr. Am. 58: 104. 1994, as “*reticulatus*”, ***emend.*** – Holotype: UNITED STATES, north-central Oregon, Nut Beds flora, Clarno Formation, UF6500 (Florida Museum of Natural History, University of Florida, United States; Manchester, 1994, pl. 54, figs. 6-13; **Fig. 6G**).

*Original species diagnosis.* – Seed reniform, bilaterally symmetrical, laterally compressed in the plane of symmetry, more or less rounded in the lateral profile, with a concave notch at the hilar end, micropyle pointed, situated adjacent to hilum; seed height (hilum to dorsal edge) 2.2-3.1, avg. 2. 7 mm (SD∼0.28, n∼ 19), width 2.2-3.0, avg. 2.5 mm (SD∼0.23, n∼l9), thickness 1.1-1.9, avg. 1.3 mm (SD∼0.20, n∼ 19); smoothly contoured, surface covered by a fine reticulum of pentagonal and hexagonal cells 49-62 um in diameter with thick anticlinal walls, these cells aligned in rows radiating from the micropyle area (Manchester, 1994).

*Emended species diagnosis.* – Differing from any other known solanaceous seeds in the character combination of seeds globose, slightly compressed, 1.3 (1.1-1.9) mm thick, 2.58 (1.16-3.27) mm long, 2.29 (0.81-2.99) mm wide; hilum lateral/sublateral, with a concave cavity at the hilar-chalazal end, and exotestal cells uniform in shape, pentagonal to hexagonal, with anticlinal walls straight on the entire seed surface.

*Species description.* – Seeds globose, slightly compressed, 1.3 (1.1-1.9) mm thick, 2.58 (1.16-3.27) mm long, 2.29 (0.81-2.99) mm wide, ratio length/width 1.15 (1.01-1.43). Hilum lateral/sublateral, with a concave cavity at the hilar-chalazal end. Exotestal cells uniform in shape, pentagonal to hexagonal, area of 0.007 (0.002-0.009) mm^2^, perimeter of 0.32 (0.2-0.51) mm. Exotestal anticlinal walls straight on the entire seed surface (**Fig. 6G**). Embryo shape unknown.

*Additional material analyzed.* – UNITED STATES, north-central Oregon, Nut Beds flora, Clarno Formation, UF9380, UF9378, UF9777, UF9382, UF9376, UF9381 (Florida Museum of Natural History, University of Florida, United States); GREECE: Rhodes, Kallithea, MB.Pb.2000/16 (Museum für Naturkunde Berlin, Germany).

*Age and distribution.* – Middle Eocene and Pleistocene, 47.8-0.01 Ma (including Manchester, 1994; Mai & Velitzelos, 2007). Complete stratigraphic range and map showing localities on **Fig. 4**.

*Remarks.* – This species comprises all fossil seed taxa clustering in cluster O of the NMDS analysis (**Fig. 2**), which includes all six analyzed specimens of *Nephrosemen* along with one extant species of *Trozellia* and one additional fossil seed previously identified as *Hyoscyamus reticulatum* (Mai & Velitzelos, 2007). The inclusion of this specimen slightly expands the circumscription of *N. reticulatum*.

Manchester (1994) suggested that this fossil might belong to either Theaceae or Solanaceae. However, the combination of a reniform seed shape, a lateral to sublateral hilum, and cerebelloid exotestal anticlinal walls provides clear support for its placement within the Solanaceae. Although this species was originally described as laterally compressed (Manchester, 1994), reexamination of the type material reveals a seed with substantial three-dimensional volume and only a slight lateral flattening. This sets *N. reticulatum* apart from strictly compressed taxa such as *Hyoscyamus undulatus* (**Fig. 5E–G**), and aligns it more closely in form with *Albionites arnensis*, the only other Solanaceae fossil taxon known to lack strong lateral compression. Nevertheless, *N. reticulatum* remains distinct from *A. arnensis* based on exotestal cell architecture. In *N. reticulatum*, the exotesta is composed of pentagonal to hexagonal cells in surface view with straight anticlinal walls, contrasting with the digitated, cerebelloid-walled cells observed in *A. arnensis*. These differences support the continued recognition of *Nephrosemen* as a morphologically distinct genus within the Solanaceae fossil record.

***Capsicum pliocenicum*** Deanna & S.D.Sm., **sp. nov.** – Holotype: ITALY, Bucine, MRSN-P/345-CCN6065b (CENOFITA collection, managed by the Regional Museum of Natural Sciences of Turin, Turin, Italy; **Fig. 7A**).

**Figure 7.**
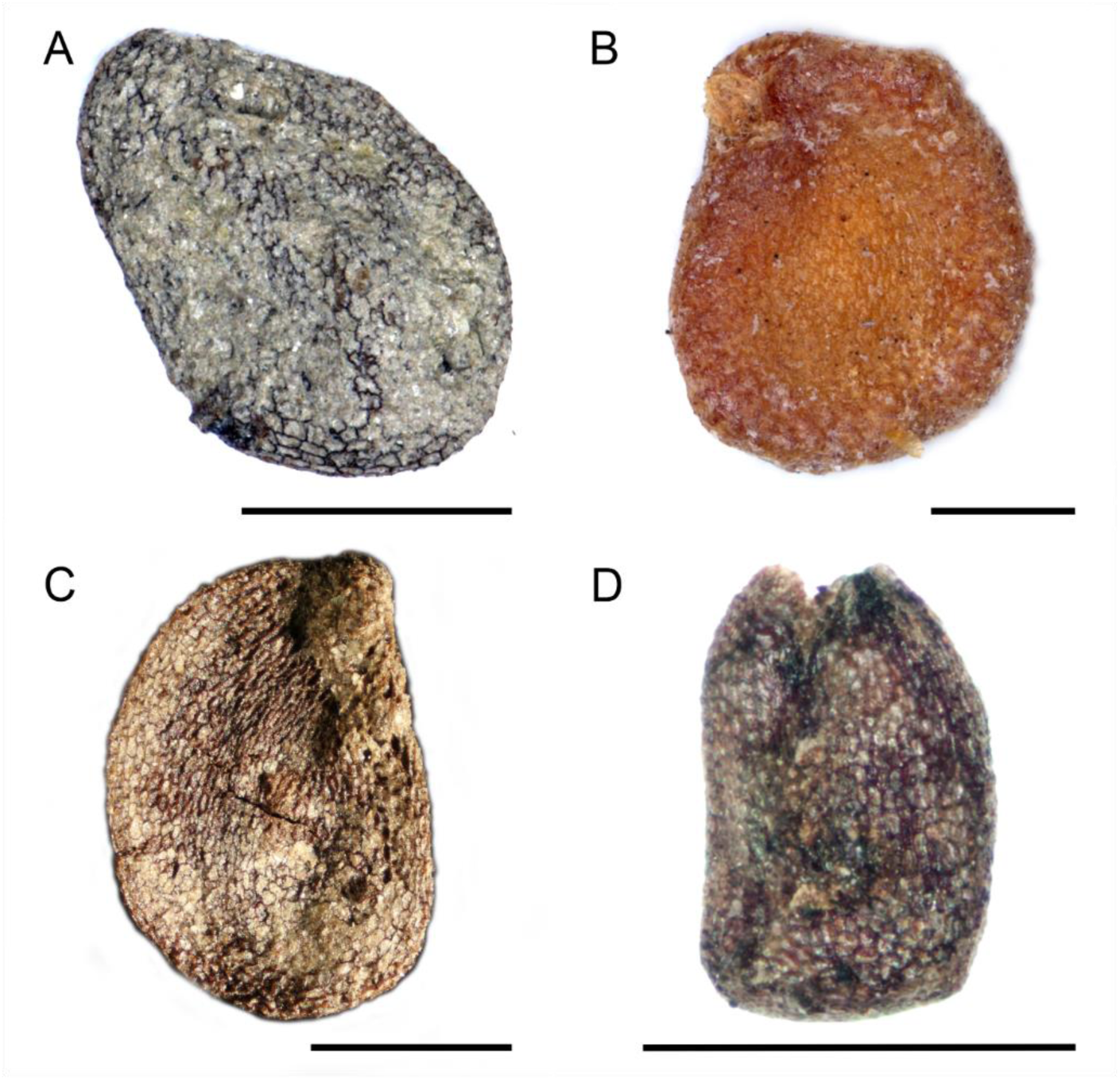
Fossils of *Capsicum pliocenicum* Deanna & S.D.Sm. sp. nov. and *Thanatosperma minutum* Deanna & S.Knapp gen. et sp. nov., and morphologically similar extant taxa. **A-B.** Specimens grouped in cluster P, *Capsicum pliocenicum*. **A.** Holotype of *C. pliocenicum*, MRSN-P/345-CCN6065b. **B.** Seed of the extant *Capsicum baccatum*. **C-D.** Representative specimens of cluster Q, *Thanatosperma minutum*. **C.** Holotype of *T. minutum*, MB.Pb.2003/0073. **D.** Paratype of *T. minutum*, BIN H676/80a. A-C photographs by R. Deanna, D by A. V. Hvalj. Bars: 1 mm.

*Species diagnosis.* – Differing from any other known solanaceous seeds in the character combination of seeds compressed, round to ovoid, hilum lateral/sublateral, without a cavity at the hilar-chalazal end, and exotestal cells varying in shape, irregularly shaped with slightly sinuate anticlinal walls in the center of the seed surface, rectangular with straight anticlinal walls towards the margins.

*Species description.* – Seeds compressed, round to ovoid, 1.71 (1.53-1.81) mm long, 1.56 (1.42-1.8) mm wide, ratio length/width 1.11 (1-1.29). Hilum lateral/sublateral, without a cavity at the hilar-chalazal end. Exotestal cells non-uniform in shape, irregular in the center, rectangular towards the margins.

Exotestal cells area of 0.004 (0.002-0.005) mm^2^, perimeter of 0.25 (0.21.0.28) mm at the center of the seed. Exotestal anticlinal walls straight on most of the seed surface, slightly sinuate in the center (**Fig. 7A**). Embryo shape unknown.

*Age and distribution.* – Pliocene to Pleistocene, 5.33-0.13 Ma (including Reid & Reid, 1907, 1915).

A complete stratigraphic range and a map showing localities are shown on **Fig. 4**.

*Additional material analyzed.* – THE NETHERLANDS, Limburg, Reuver, RGM 792900; THE NETHERLANDS, Limburg, Tegelen, RGM 793332 (Naturalis Biodiversity Center, Leiden, The Netherlands).

*Etymology.* – The name of this species relates to its stratigraphic distribution, with most of its fossil specimens assigned to the Pliocene.

*Remarks.* –This species comprises all fossil seed taxa clustering in cluster P of the NMDS analysis (**Fig. 2**), which includes only three fossils and one extant species of the genus *Capsicum.* Despite their small size, smaller even than the seeds of extant *Capsicum* species, the fossils share a distinctive combination of characters that support their assignment to this genus. Key diagnostic features include a lateral hilum position, absence of a hilar-chalazal cavity, and predominantly straight exotestal anticlinal walls.

The fossil material includes specimens previously identified as *Solanum* (Reid & Reid, 1907, 1915) due to superficial similarities in seed shape and size. However, our refined micromorphological assessment and NMDS analysis reveal closer affinities with *Capsicum* L. These fossils originate from Pliocene and Pleistocene deposits in the Netherlands and Italy, marking one of the earliest known occurrences of *Capsicum*-like seeds in Europe.

***Thanatosperma*** Deanna & S.Knapp., **gen. nov. –** Type: *Thanatosperma minutum*

*Generic diagnosis.* **–** Differing from any other known solanaceous seeds in the character combination of seeds compressed, hilum terminal, without a cavity at the hilar-chalazal end, and exotestal cells uniform, rectangular to hexagonal, with straight anticlinal walls on the entire seed surface.

*Etymology.* **–** The name is a combination of the words *thanatos* - meaning death, and *sperma* - meaning seed, in reference to the extinct nature of this taxon.

***Thanatosperma minutum*** Deanna & S.Knapp, **sp. nov.** – Holotype: GERMANY, Oberzella a. d. Werra, MB.Pb.2003/0073 (Museum für Naturkunde Berlin, Germany; **Fig. 7C**).

*Species diagnosis*. – Same as for genus.

*Specific description*. – Seeds compressed, ovoid to oblong, 1.67 (1.07-2.19) mm long, 1.16 (0.68- 1.54) mm wide, ratio length/width 1.47 (1.23-1.67). Hilum terminal, slightly protruding, without a cavity at the hilar-chalazal end. Exotestal cells uniform, rectangular to hexagonal, area of 0.004 (0.001-0.005) mm^2^, perimeter of 0.23 (0.14-0.3) mm. Exotestal anticlinal walls straight on the entire seed surface (**Fig. 7C-D**). Embryo shape unknown.

*Age and distribution.* – Pliocene to Holocene, 5.33-0.001 Ma (including Carvallo & Martinetto, 2001; Girotti & al., 2003; Gümbel & Mai, 2004). A complete stratigraphic range and a map showing localities are shown on **Fig. 4**.

*Additional material analyzed.* – ITALY, Castelletto Cervo II, MRSN-P/345-CCN 448; ITALY, Torre Picchio, MRSN-P/345-CCN 3067 (CENOFITA collection, managed by the Regional Museum of Natural Sciences of Turin, Turin, Italy); RUSSIA, West Siberia, Novosibirsk oblast, 55.166345, 82.857844, BIN H676/80a (Komarov Botanical Institute, St. Petersburg, Russia).

*Etymology.* – The specific epithet relates to the small length of the seeds.

*Remarks.* – This species comprises all fossil seed taxa clustering in cluster Q of the NMDS analysis (**Fig. 2**), including only four seed fossils. Although some of these fossils were previously attributed to *Solanum* or *Hyoscyamus* (Carvallo & Martinetto, 2001; Girotti & al., 2003; Gümbel & Mai, 2004), their exclusion from any extant taxonomic grouping in the NMDS analysis (**Fig. 2**) supports their interpretation as belonging to an extinct genus.

The combination of an ovoid to oblong, laterally compressed seed shape, a terminal hilum, and straight exotestal anticlinal walls sets this taxon apart from all other known fossil and extant Solanaceae seeds. This unique morphological profile suggests it represents a distinct evolutionary lineage within the family that is no longer found in the modern flora.

## DISCUSSION

Our paper presents the first comprehensive analysis of the Solanaceae seed fossil record. By reviewing taxonomic literature and analyzing 110 fossil specimens together with 354 living species, we created a new classification scheme that recognizes six new species and four new genera. Our treatment retains two previously described genera and three species but uses additional specimens to revise and expand their descriptions. Using this new taxonomic framework, we find that several seed fossil species trace their history to the Eocene (ca. 48 Ma) while the greatest diversity of seed fossil taxa is found during the Neogene (ca. 23-2.5 Ma; **Fig. 4**). All of these seed fossils were recovered from the Northern Hemisphere and are spread from the Americas to Eurasia, supporting the emerging view that the wide geographic distribution of the Solanaceae was achieved at least by the Eocene.

### Morphological diversity of nightshade seed fossils

Our multivariate analysis using both discrete and continuous traits revealed that morphology of seed fossils span much of the morphological diversity found in extant taxa. These include fossil taxa like *Solanum foveolatum* and *Capsicum pliocenicum* with the flattened, round or reniform seeds common in extant berry clade species, as well as fossil taxa with globose seeds (*Albionites arnensis*, *Nephrosemen reticulatum*). These fossil seeds also varied in size from less than 1 mm in length to over 3 mm, similar to the range of most extant Solanaceae seeds (1.2 to 2.5 mm; Hunziker, 2001) although these fossils do not reach the larger end of the spectrum (e.g., 6.5 mm long in *Solandra* Sw., 30 mm in *Duckeodendron*). Consistent with the taxonomic utility of seed coat characters in previous taxonomic studies (e.g., *Schwenckia* L., Carvalho & al., 1999; *Physalis*, Zhang & Wen, 1996; Hyoscyameae, Zhang & al., 2005), we also found significant variation in the size and shape to the exotestal cells and their anticlinal walls, from the tiny uniform straight-walled cells of *N. reticulatum* (**Fig. 6G**) to the variable sinuate-cerebelloid cells of *Solanoides dorofeevii* (**Fig. 6A-C**). Finally, our micro-CT scans confirmed that the fossil seed record includes curved (*S. dorofeevii*; **Fig. 1D**) and coiled embryos (BIN K453/175; Dorofeev, 1966; **Fig. 1C**).

Although all our fossil descriptions belonged to well-defined clusters within extant Solanaceae seed morphospace (**Fig. 2**), two of the ten recognized taxa did not group tightly with any extant taxa. *Solanispermum reniforme*, represented by nine specimens, has some common nightshade seed traits (flattened reniform shape and a distinct hilar cavity, **Fig. 5C-D**), but these appear in combination with less common traits (mostly straight exotestal anticlinal walls and a spongy fibrous coat). *Solanum miocenicum*, also represented by nine fossils, has flattened round seeds with the common lateral hilum but lacks a hilum cavity (**Fig. 5A-B**). Nevertheless, as some *Solanum* species also lack this cavity and previous authors assigned this material to *Solanum* (Reid & Reid, 1915; Dorofeev, 1963), we retained this species, provisionally, in the extant genus. Overall, we chose to conservatively assign fossil specimens to fossil taxa, unlike previous authors (Dorofeev 1960, 1977; Mai & al., 1963; Basilici & al., 1997; Mai, 2010).

In comparing these fossil seeds with extant taxa, it is important to note aspects of fossilization that may influence their morphology and diversity and, in turn, our conclusions about their taxonomic affinities. First, all biological material experiences some degree of compression during fossilization, suggesting that we should be cautious about relying too strongly on lateral compression as a taxonomic character. In our analysis, compression was among the strongest characters separating taxa, although discrete morphological traits (exotestal cell wall shape, hilum position, presence of a hilar cavity) were similarly, and in some cases, more informative (Table S6). Also, we recovered two fossil taxa without lateral compression (*N. reticulatum* and *A. arnensis*; **Fig. 2**; Table S3). Thus, while the seeds analyzed here likely experienced some compression during fossilization, our sample retains three-dimensional variation that mirrors the range in extant taxa, from flattened to globose, adding to our ability to delimit distinct clusters. While these fossil seeds span much of the seed shape diversity within the family, the notable lack of large fossil seeds may simply be also attributable to the process of fossilization, as larger material is often washed away, leaving the small seeds (typically <1cm) comprising the bulk of paleocarpological material. Combined with the rarity of larger seeds in the family in general, it is not surprising to be missing large seeds from the fossil record.

### Evolutionary and biogeographic implications of stratigraphic history

While Solanaceae fruit fossils have only been recovered from the Americas, seed fossils indicate a broader geographic distribution, spanning North America, Europe and Asia (**Fig. 4**). Indeed, two of the three earliest seed fossils (*Solanispermum reniforme*, 48 Ma; *Albionites arnensis*, 48 Ma) were recovered from Europe and the other (*Nephrosemen reticulatum*, 47.8 Ma) from North America. Their ages are similar to the fruit fossils (also mid to late Eocene, spanning 52-34 Ma; Deanna & al., 2023). However, unlike the fossil fruit history, the Solanaceae seed fossil record extends entirely toward the present, thanks to the large amount of material available to estimate the stratigraphic range (**Fig. 4**). Two of the Eocene seed fossil taxa (*S. reniforme* and *N. reticulatum*) have stratigraphic ranges reaching the Pleistocene as do all of the taxa that first appeared in the Oligocene. With the persistence of these Eocene and Oligocene fossils together with new taxa that appear in the Miocene (*Hyoscyamus undulatus*) and Pliocene (*Capsicum pliocenicum*, *Thanatosperma minutum*), the taxonomic richness is highest at the Pliocene/Pleistocene boundary (c. 2.58 Ma; **Fig. 4**). As the majority of these fossil seeds present a flattened form, which is almost exclusively found in solanaceous berries (Campos, 2023), this Neogene seed diversity likely resulted from the radiation of major berry-bearing lineages of Solanaceae (Solaneae, Capsiceae, Physalideae; Deanna & al., 2023; Zhang & al., 2023).

Although these flattened seeds typical of those in extant berry fruits are the most abundant form in the fossil record, it is notable that the two taxa without compression (*Albionites arnensis* and *Nephrosemen reticulatum*) are also among the oldest (**Fig. 4**). Such small globose seeds are often associated with capsules (e.g., in the extant genera *Schizanthus* and *Petunia*), and this fruit type, shared with the sister family Convolulaceae, has been inferred as the ancestral state (Knapp, 2002; Pabón-Mora & Litt, 2011). While our fossil stratigraphy demonstrates the early appearance of globose solanaceous seeds, neither of these fossils could be confidently placed within any extant genus. The lack of any seeds or fruits clearly belonging to any of the diverse capsular clades (Schwenckieae, Petunieae, Nicotianoideae) remains a glaring gap in the fossil record. Thus, incorporating fossil information into the estimation of the ancestral state for fruits (as in Finarelli & Flynn, 2006) awaits additional taxa with more definitive diagnostic traits.

Our study also reports the first case of a fossilized elaiosome, a specialized food body that is tightly linked with ant dispersal (Gorb & Gorb, 2003). Elaiosomes provide a nutrient-dense food source and also contain compounds that induce carrying behavior (Brew & al., 1989), prompting ants to transport the seed to their underground nests. After consuming the elaiosome, ants typically remove the bare seed from the nest, and through this multistage transportation process, facilitate seed dispersal (Servigne & Detrain, 2010). In *Datura*, the large elaisome forms from seed tissue just above the funicle, and *Hyoscyosperma daturoides* displays a similar elaiosome position near the hilum (**Figs. 1A-B**, **3A**). Importantly, the elaiosome tissue in *H. daturoides* connects through the seed coat, confirming its development from seed tissue (Suppl. Video in OSF link provided after revisions). This fossil first appears ca. 27.8 Ma, although angiosperm-wide surveys suggest that myrmecochory may have arisen as early as 100 Ma (Lengyel & al., 2010), shortly after the first appearance of ants in the fossil record (Lepeco & al., 2025). Hence it appears that at least since the Oligocene, Solanaceae have relied upon ants in addition to frugivores and abiotic forces for seed dispersal.

These diverse dispersal mechanisms help to explain the wide distribution of the family, although the biogeographic origins of the family remain unclear. With its concentration of diversity in the Andes, the Solanaceae has long been considered to have originated in South America (Olmstead 2013; Dupin & al., 2017). While the oldest fossils of any type are the *Physalis* fruit from Laguna del Hunco (52 Ma) in Argentina (Wilf & al., 2017; Deanna & al., 2020), the chili pepper clade fossil from Colorado (*Lycianthoides calycina* Deanna & Manchester, c. 50 Ma; Deanna & al., 2023) and several of the Northern Hemisphere seeds described here are of similar age. Their occurrence coincides with the Early Eocene Climatic Optimum (EECO, 53.3 to 49.1 Ma), a period of warm temperatures, high precipitation and high carbon dioxide levels (Zachos & al., 2001, 2008; Cramwinckel & al., 2023) with corresponding high plant diversity across the Southern and Northern hemispheres (Wilf & al., 2003; Woodburne & al., 2009). Plant families with similarly wide distributions by the EECO, such as the Asteraceae (Mandel & al., 2019) and Scrophulariaceae (Villaverde & al., 2023), trace their origins to the late Cretaceous (100 to 68 Ma), which may also be the case for Solanaceae. While a South American origin followed by dispersal to the north and east is consistent with the fossil record (**Fig. 4**), it is intriguing that the oldest fossils (58 Ma) of the sister family (Convolvulaceae) are from India (Srivastava & al., 2018), opening the possibility for dispersal in the opposite direction with later colonization and diversification in South America.

## Conclusions

Understanding the timeline of Solanaceae diversification has long been hampered by the paucity of fossils and uncertainty in their assignment. Our work demonstrates that seeds provide a wealth of morphological variation that can form the basis for a robust classification, with more recent fossils presenting clade-specific traits that allow them to be confidently assigned to particular genera. This assignment of the large pool of fossil seeds to individual taxa gives clear direction for future studies to address taxonomic and biogeographic gaps. In particular, the Solanaceae fossil record (across all seeds, fruit, pollen, and wood) entirely lacks representation from Africa and Oceania, despite the many distinct lineages (Nicotianoideae, Hyoscyameae, *Mandragora* L.) with high diversity in those regions. Moreover, the seed fossil record does not capture the full range of extant seed sizes and morphologies (e.g., the presence of wings), as many of these extremes fall into less diverse groups with a correspondingly lower chance of being preserved in the fossil record. Nevertheless, the morphological variation found among the seeds described paints a clear picture of the burst of diversity at the Paleogene / Neogene boundary and lays out a multivariate morphometric approach that will expedite the description of unexplored collections by comparison with seeds from living and extinct species. In combination with the new records of Solanaceae fruit fossils, these fossil seeds will open new doors for calibrating the family tree and addressing the outstanding questions about the evolutionary backdrop for nightshade diversification.

## AUTHOR CONTRIBUTIONS

R. Deanna led the data collection, conducted the analyses, interpreted the results, designed the figures, and prepared the taxonomic descriptions. A. V. Hvalj, E. Martinetto, E.-M. Sadowski, and S. Manchester contributed to the analysis and interpretation of the fossil specimens and provided paleobotanical training to R. Deanna. A. V. Hvalj and E. Martinetto also photographed the fossil material. V. Fernandez performed the micro-CT scans and trained R. Deanna in image processing and data interpretation. S. Knapp, A. Campos, G. E. Barboza, E. Dean, T. Särkinen, F. Chiarini, G. Bernardello, R. Deanna, and S. D. Smith scored morphological characters of extant Solanaceae species. H. Sauquet provided access to and support with the PROTEUS database. R. Deanna and S. D. Smith designed the study and wrote the manuscript, with significant input and revisions from all co-authors.

## ACKNOWLEDGEMENTS

This work was supported by the National Science Foundation (DEB 1902797 to SDS) and a grant to RD from the SYNTHESYS+ project (grant agreement 823827, financed by the European Community Research Infrastructure Actions under H2020-INFRAIA-2018-2020; Integrating and opening research infrastructures of European interest). We are grateful to Thomas Denk (Swedish Museum of Natural History, Stockholm) for valuable discussions on the evolutionary history of Tertiary floras. We thank Vladimir Dorofeyev, curator of the herbarium of higher plants at the Komarov Botanical Institute (LE), and Veronika Romanova and Kseniya Naumova, curators of the carpological collections at the Komarov Botanical Museum, for facilitating access to important historical seed material. We are especially indebted to the staff from the Imaging and Analysis Centre (IAC) at the Natural History Museum, London, and to the NHM collections management staff for their generous support and for granting access during pandemic-related lockdowns. We also thank staff from all the herbaria and fossil collections cited in the Methods section for their assistance and access to specimens. Catrin Puffert, Cornelia Hiller (Museum für Naturkunde, Berlin), and Sofie De Smedt (Meise Botanical Garden) provided access to literature and specimens in the Palaeobotany collection and offered technical support throughout our visits. We are also grateful to Adrian Carper (University of Colorado, Boulder) for his support with microscopy and imaging, and to Silvana Mosteschiesi (CORD) for creating the maps presented in Figure 4. RD also acknowledges the financial support from the Consejo Nacional de Investigaciones Científicas y Técnicas (CONICET, Argentina), the Secretaría de Ciencia y Tecnología de la Universidad Nacional de Córdoba (grant 203/14, SECYT-UNC, Argentina), and the European Union under the Marie Skłodowska-Curie grant agreement No 101151612 (MSCA).

## LEGENDS - SUPPLEMENTARY FIGURES

**Figure S1.**
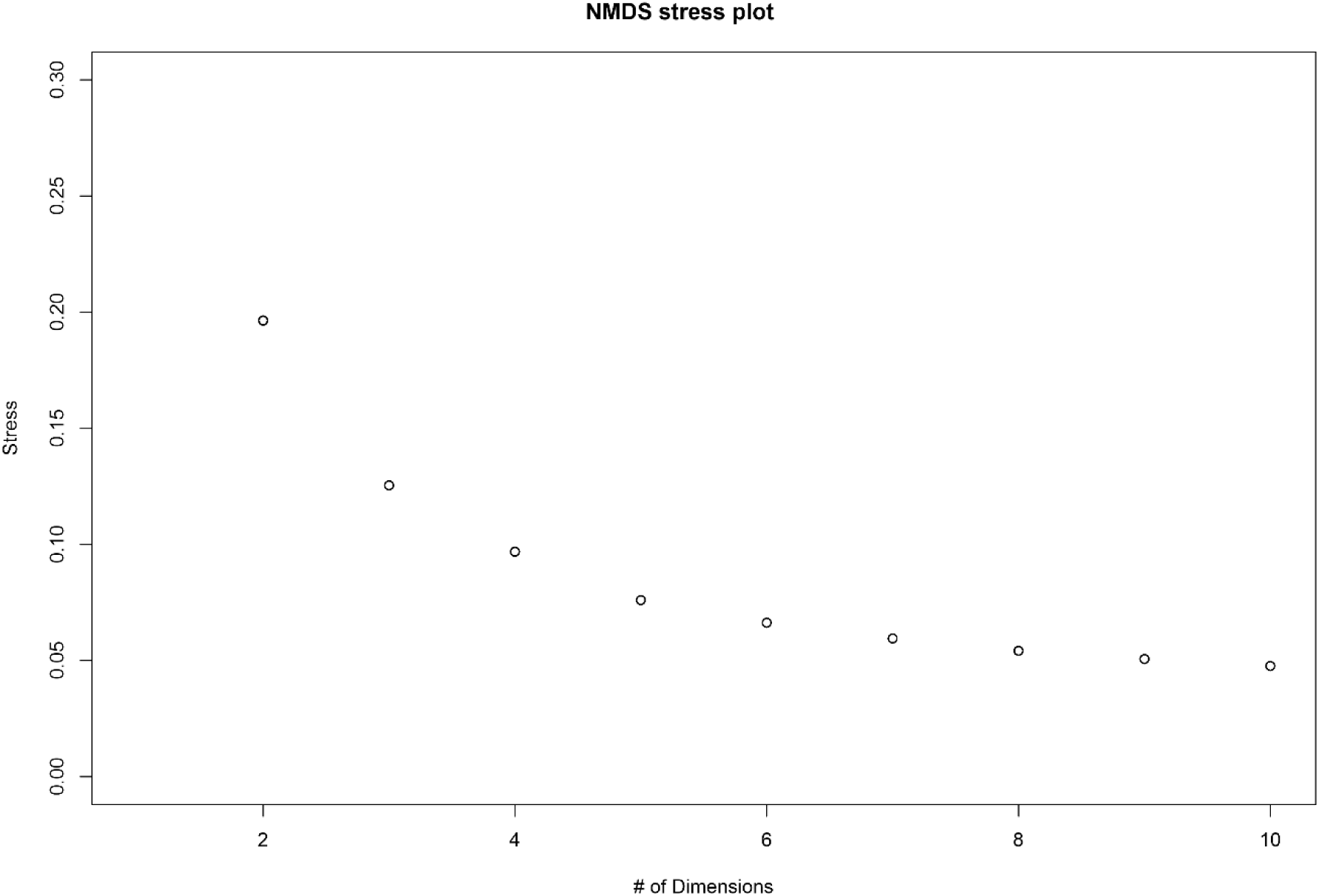
Stress plot for non-metric multidimensional scaling (NMDS) analysis. The plot shows the relationship between the number of dimensions and the stress value, used to assess the optimal number of dimensions for representing the seed morphometric data. Lower stress values indicate a better fit; a stress value below 0.2 is generally considered acceptable.

## LEGENDS - SUPPLEMENTARY TABLES

**Table S1.** All published and unpublished claims for Solanaceae seeds in the fossil record and extant taxa according to the clusters of Fig. 2. Specimens with an asterisk were analyzed using a high-resolution Zeiss Versa 520 micro-CT system to determine their systematic affinity in Särkinen et al. (2018; *Solanispermum reniforme*) or this publication. The geological time scale v. 2024/12 was used (Dec 2024; updated). Na means no available information. All cited references appear in the main text.

**Table S2.** Seed traits scored for fossil specimens and extant species of Solanaceae. Bold characters are secondary traits, calculated from primary measurements; italicized characters were not included in the NMDS analysis. Characters marked with an asterisk (*) were scored using high-resolution micro-CT. Roundness compares the shape of an object to a perfect circle, with a value of 1 indicating a perfect circle and values closer to 0 representing more elongated or irregular shapes.

**Table S3.** Morphometric data for fossil specimens and extant taxa. All measurements are given in millimeters. See Table S2 for coding scheme for discrete characters. The raw data tab contains all the original measurements, including multiple measurements for exotestal wall area, perimeter, and roundness; the clean data tab presents the data that was directly input into the multivariate analyses including, averages when multiple measurements were taken. Fossil specimens are grouped and labeled by cluster from the NMDS analysis in the clean data tab.

**Table S4.** Voucher information and bibliographic references for all extant taxa used in the study, including the sources for each scored character.

**Table S5.** Micro-CT scanning parameters and settings for fossil and extant Solanaceae seeds. All specimens were imaged using a Zeiss Xradia 520 Versa system, with settings modified for optimal visualization. Bolded species names indicate fossil taxa, and asterisks indicate that they are described here.

**Table S6.** Discrete and continuous characters analyzed, including scores that show the contribution of each variable to each axis of the NMDS and significance. The coding scheme for each character is shown in Table S2. Significance codes: 0 ‘***’ 0.001 ‘**’ 0.01 ‘*’ 0.05 ‘.’ 0.1 ‘ ’ 1.

