## Supplementary table S1 for "Seed fossil record of Solanaceae revisited"

**Table S1.** All published and unpublished claims for Solanaceae seeds in the fossil record and extant taxa according to the clusters of Fig. 2. Specimens with an asterisk were analyzed using a high-resolution Zeiss Versa 520 micro-CT system to determine their systematic affinity in Särkinen et al. (2018; *Solanispermum reniforme*) or this publication. The geological time scale v. 2024/12 was used (Dec 2024; updated). Na means no available information. All cited references appear in the main text or Table S4.

| Specimen code / *extant species* -identification | Reference (or unpublished) / Voucher or barcode | Epoch: Stage (age) | Country and Occurrence (coordinates if available) | Repository / Herbarium acronym |
| --- | --- | --- | --- | --- |
| **CLUSTER A – HYOSCYOSPERMA DATUROIDES** | | | | |
| K453/133* – Holotype (designated here) –identified as *Physalis pliocenica* **(Fig. 1A-B, 3A)** | Dorofeev (1957a) | Pliocene (5.33 -2.58 Ma) | 47.762692, 42.746863, Rostov oblast, Northern Caucasus, Russia. | Komarov Botanical Institute, St. Petersburg, Russia. |
| H1594/44 | Unpublished | Late Oligocene (27.8-23 Ma) | Tomsk oblast, West Siberia, Russia. | Komarov Botanical Institute, St. Petersburg, Russia. |
| H4044/71* (**Fig. 3C)** | Unpublished | Middle Miocene (16-11.6 Ma) | 54.30109, 68.346153, Soltústik Qazaqstan, Kazakhstan. | Komarov Botanical Institute, St. Petersburg, Russia. |
| H554 | Unpublished | Quaternary (2.58-0 Ma) | 55.903982, 74.727913, Omsk oblast, West Siberia, Russia. | Komarov Botanical Institute, St. Petersburg, Russia. |
| *Hyoscyamus niger* L.* (**Fig. 3D)** | BM001121613 | Current (0 Ma) | Na | Natural History Museum, London, UK. |
| *Datura metel* L.  **(Fig. 3B)** | KIB7415 | Current (0 Ma) | Na | Komarov Botanical Institute, St. Petersburg, Russia. |
| **CLUSTER B – not treated (bibliography only)** | | | | |
| KRAM-P 242/163 | Velichkevich & Zasfawniak (2003) | Pliocene (5.33 -2.58 Ma) | Valleys of rivers Dnieper, Pripyat’, Nemen, Belarus. | Institute of Geological Sciences, the National Academy of Sciences of Belarus in Minsk, Belarus. |
| MINM-P-BGO-2/162 | Velichkevich & Zasfawniak (2003) | Pliocene (5.33 -2.58 Ma) | Valleys of rivers Dnieper, Pripyat’, Nemen, Belarus. | Institute of Geological Sciences, the National Academy of Sciences of Belarus in Minsk, Belarus. |
| **CLUSTER C – SOLANUM FOVEOLATUM** | | | | |
| MB.Pb.1998/0434 – Holotype *Solanum foveolatum* Negru (**Fig. 3E**) | Negru (1986) | Early Miocene (23.03-13.82 Ma) | Bauersberg, Germany. | Museum für Naturkunde Berlin, Germany. |
| K390* – identified as *Physalis alkekengi* L. (currently *Alkekengi officinarum* Moench) | Dorofeev (1960a) | Late Miocene / Early Pliocene (11.63-3.6 Ma) | 54.834631, 56.397617, Bashkiria, Ural, Russia. | Komarov Botanical Institute, St. Petersburg, Russia. |
| K587B – identified as *Physalis alkekengi* **(Fig. 3F;** currently *Alkekengi officinarum* Moench) | Dorofeev (1977) | Late Miocene / Early Pliocene (11.63-3.6 Ma) | 54.753123, 54.883264, Bashkiria, Ural, Russia. | Komarov Botanical Institute, St. Petersburg, Russia. |
| H4759 | Unpublished | Late Oligocene (27.82-23.03 Ma) | 58.328408, 79.403543, Tomsk oblast, West Siberia, Russia. | Komarov Botanical Institute, St. Petersburg, Russia. |
| K690 | Unpublished | Late Pliocene (3.6-2.58 Ma) | 52.158587, 30.623945, Kholmyech, Belarus. | Komarov Botanical Institute, St. Petersburg, Russia. |
| K536 | Unpublished | Oligocene (33.9-23.03 Ma) | 60.100319, 83.209543, Tomsk oblast, West Siberia, Russia. | Komarov Botanical Institute, St. Petersburg, Russia. |
| H4308 | Unpublished | Early Oligocene (33.9-27.82 Ma) | 54.952243, 82.814527, Novosibirsk oblast, West Siberia, Russia. | Komarov Botanical Institute, St. Petersburg, Russia. |
| H1818 | Unpublished | Early Miocene (23.03-13.82 Ma) | 55.266820, 79.146436, Novosibirsk oblast, West Siberia, Russia. | Komarov Botanical Institute, St. Petersburg, Russia. |
| H4152 | Unpublished | Late Oligocene (27.82-23.03 Ma) | 60.108504, 83.242754, Tomsk oblast, West Siberia, Russia. | Komarov Botanical Institute, St. Petersburg, Russia. |
| H1328 | Unpublished | Late Oligocene (27.82-23.03 Ma) | 57.021166, 82.071869, Tomskaya oblast, West Siberia, Russia. | Komarov Botanical Institute, St. Petersburg, Russia. |
| H1120 | Unpublished | Late Oligocene (27.82-23.03 Ma) | 56.726710, 86.453605, Tomsk oblast, West Siberia, Russia. | Komarov Botanical Institute, St. Petersburg, Russia. |
| H3564 | Unpublished | Early Miocene (23.03-13.82 Ma) | 56.648865, 73.015980, Omsk oblast, West Siberia, Russia. | Komarov Botanical Institute, St. Petersburg, Russia. |
| MB.Pb.1993/7605 – identified as *Physalis pliocenica* | Mai (2000) | Early Miocene (23.03-13.82 Ma) | Hartau b. Zittau, Germany. | Museum für Naturkunde, Berlin, Germany. |
| MB.Pb.1993/9655B | Unpublished | Pleistocene (2.58-0 Ma) | Mahlis, Germany. | Museum für Naturkunde, Berlin, Germany. |
| MB.Pb.1993/6831* | Unpublished | Late Pliocene (3.6-2.58 Ma) | Nordhausen, Germany. | Museum für Naturkunde, Berlin, Germany. |
| MB.Pb.1993-5546* | Unpublished | Late Pliocene (3.6-2.58 Ma) | Berga b. Sangerhausen, Germany. | Museum für Naturkunde, Berlin, Germany. |
| MB.Pb.1993/2220 – identified as *Physalis pliocenica* | Mai (1997) | Late Oligocene (27.82-23.03 Ma) | Horka, Germany. | Museum für Naturkunde, Berlin, Germany. |
| MB.Pb.1993/10814 – identified as *Solanum dulcamara* L. | Mai (2010) | Pleistocene (2.58-0 Ma) | Braunsbedra b. Merseburg, Germany. | Museum für Naturkunde, Berlin, Germany. |
| MB.Pb.1993/0791 – identified as *Solanum dulcamara* L. **(Fig. 3G**) | Mai & al. (1963) | Late Pliocene (3.6-2.58 Ma) | Rippersroda b. Arnstadt, Germany. | Museum für Naturkunde, Berlin, Germany. |
| MRSN-P/345-CCN4332 – identified as *Solanum* cf. *dulcamara* | Basilici & al. (1997) | Pliocene: Zanclean (5.33-3.6 Ma) | Sento-S7a, Italy. | CENOFITA collection, managed by the Regional Museum of Natural Sciences of Turin, Turin, Italy. |
| *Cuatresia garciae* Hunz. | Martinez 1410 (NY) | Current (0 Ma) | Na | New York Botanical Garden, New York, USA. |
| *Anisodus luridus* Link ex Spreng. (**Fig. 3H**) | KIB7393 | Current (0 Ma) | Na | Komarov Botanical Institute, St. Petersburg, Russia. |
| **CLUSTER D – not treated (only broken seed)** | | | | |
| K453/175* – identified as *Physalis alkekengi* (currently *Alkekengi officinarum* Moench; **Fig. 1C**) | Dorofeev (1966) | Pliocene (5.33 -2.58 Ma) | 47.762692, 42.746863, Rostov oblast, Northern Caucasus, Russia. | Komarov Botanical Institute, St. Petersburg, Russia. |
| *Solanum melongena* L. | Weber s.n. (FLAS) | Current (0 Ma) | Na | FLAS |
| *Solanum citruliifolium* A.Braun | Weber s.n. (FLAS) | Current (0 Ma) | Na | FLAS |
| *Physalis angulata* L.* | BM000941947 | Current (0 Ma) | Na | BM |
| *Solanum sisymbriifolium* Lam. | Weber s.n. (FLAS) | Current (0 Ma) | Na | FLAS |
| *Solanum pyracanthos* Lam. | Weber s.n. (FLAS) | Current (0 Ma) | Na | FLAS |
| *Solanum carolinense* L. | Pharmacy Dept. s.n. (FLAS) | Current (0 Ma) | Na | FLAS |
| *Physochlaina physaloides* G.Don | Unknown collector 1012 (COLO) | Current (0 Ma) | Na | COLO |
| *Solanum bicolor* Willd. | Rhoads s.n. (FLAS) | Current (0 Ma) | Na | FLAS |
| *Physalis minima* L. | KIB7401 | Current (0 Ma) | Na | Komarov Botanical Institute, St. Petersburg, Russia. |
| *Solanum bahamense* L. | Solanaceae Source (2020) | Current (0 Ma) | Na | Na |
| *Solanum juvenale* Thell. | Wahlert & al. (2015) | Current (0 Ma) | Na | Na |
| *Solanum pseudocapsicum* L. | Knapp (2002) | Current (0 Ma) | Na | Na |
| *Physalis walteri* Nutt. | Majure 3051 (FLAS) | Current (0 Ma) | Na | FLAS |
| **CLUSTER E – not treated (bibliography only)** | | | | |
| Identified as *Physalis* cf. *alkekengi* (currently *Alkekengi officinarum* Moench) | Szafer (1946) | Pliocene (5.33 -2.58 Ma) | Kroscienko, Poland. | Na |
| Identified as *Physalis* cf. *alkekengi* (currently *Alkekengi officinarum* Moench) | Palamarev (1970) | Pliocene: Zanclean (5.3-3.6 Ma) | Baldevo Formation, Garmen, Bulgaria. | IB, Department of Paleobotany and Pollen Analysis at the Institute of Botany at Bulgarian Academy of Sciences (BAS), Bulgaria. |
| **CLUSTER F – not treated (bibliography only)** | | | | |
| Identified as *Hyoscyamus* sp. | Szafer (1946) | Pliocene (5.33 -2.58 Ma) | Kroscienko, Poland. | Na |
| *Solanum homalospermum* Chiarini | Chiarini & al. 505 (CORD, NY) | Current (0 Ma) | Na | CORD, NY |
| *Solanum sessile* Ruiz & Pav.* | Knapp & al. 6316 (BM) | Current (0 Ma) | Na | BM |
| *Solanum profusum* C.V. Morton | Valtuone 58 (CORD) | Current (0 Ma) | Na | CORD |
| *Quincula lobata* (Torr.) Raf. | Freeman 14549 (COLO) | Current (0 Ma) | Na | COLO |
| *Mandragora caulescens* C.V.Clarke | Banfford 29085 (NY) | Current (0 Ma) | Na | NY |
| **CLUSTER G – SOLANUM MIOCENICUM** | | | | |
| K528/50 – Holotype (designated here) – Identified as *Solanum* sp. **(Fig. 5A)** | Dorofeev (1963) | Miocene (23.03-5.33 Ma) | 57.734521, 71.173771, Omsk oblast, West Siberia, Russia. | Komarov Botanical Institute, St. Petersburg, Russia. |
| H4177/82 (**Fig. 5B)** | Unpublished | Early Miocene (23.03-15.97 Ma) | 56.671976, 74.706109, Omsk oblast, West Siberia, Russia. | Komarov Botanical Institute, St. Petersburg, Russia. |
| H2373 | Unpublished | Late Oligocene (27.82-23.03 Ma) | 53.839196, 77.443347, Novosibirsk oblast, West Siberia, Russia. | Komarov Botanical Institute, St. Petersburg, Russia. |
| K527 – Identified as *Solanum* sp. | Dorofeev (1963) | Miocene (23.03-5.33 Ma) | 57.693604, 71.393553, Omsk oblast, West Siberia, Russia. | Komarov Botanical Institute, St. Petersburg, Russia. |
| H1344 | Unpublished | Oligocene / Miocene (33.9-5.33 Ma) | 61.44233, 83.864155, Nizhnevartovsky, Janty-Mansi, West Siberia, Russia. | Komarov Botanical Institute, St. Petersburg, Russia. |
| H2000 | Unpublished | Late Oligocene (27.82-23.03 Ma) | 56.049558, 84.198969, Novosibirsk oblast, West Siberia, Russia. | Komarov Botanical Institute, St. Petersburg, Russia. |
| MB.Pb.1993/9655 | Unpublished | Pleistocene (2.58-0 Ma) | Mahlis, Germany. | Museum für Naturkunde, Berlin, Germany. |
| MB.Pb.1993/4558 | Unpublished | Late Miocene (13.82-5.33 Ma) | Bröthen b. Hoyerswerda, Germany. | Museum für Naturkunde, Berlin, Germany. |
| **CLUSTER H – SOLANISPERMUM RENIFORME** | | | | |
| GSM76684 – Holotype *Solanispermum reniforme* Chandler | Chandler (1957); Reid & Reid (1910) | Uncertain (Oligocene?) | Bovey Tracey, United Kingdom. | British Geological Survey, Keyworth, Nottinghamshire, United Kingdom. |
| MRSN-P/345-CCN6065a | Unpublished | Pleistocene (2.58-0 Ma) | Bucine, Italy. | CENOFITA collection, managed by the Regional Museum of Natural Sciences of Turin, Turin, Italy. |
| MRSN-P/345-CCN4328a | Martinetto (1995) | Pliocene: Zanclean (5.3-3.6 Ma) | Ca' Viettone, Italy. | CENOFITA collection, managed by the Regional Museum of Natural Sciences of Turin, Turin, Italy. |
| MRSN-P/345-CCN4328b | Martinetto (1995) | Pliocene: Zanclean (5.3-3.6 Ma) | Ca' Viettone, Italy. | CENOFITA collection, managed by the Regional Museum of Natural Sciences of Turin, Turin, Italy. |
| V-42019* – *Solanispermum reniforme* Chandler (paratype) | Chandler (1962) | Eocene: mid to late Lutetian (46-44 Ma) | Branksome Dene, Bournemouth Freshwater Beds, Branksome Sand Formation, United Kingdom. | Natural History Museum, London, United Kingdom. |
| V-40891* – *Solanispermum reniforme* (paratype, **Fig. 5C**) | Chandler (1962) | Eocene: late Ypresian/early Lutetian (48-46 Ma) | Arne, Poole Formation Sandbanks, Poole Formation, United Kingdom. | Natural History Museum, London, United Kingdom. |
| MB.Pb.1993/5544 – Identified as *Solanum dulcamara* L. | Mai & Walther (1988) | Late Pliocene (3.6-2.58 Ma) | Berga b. Sangerhausen, Germany. | Museum für Naturkunde, Berlin, Germany. |
| MB.Pb.1993/5544a | Unpublished | Late Pliocene (3.6-2.58 Ma) | Berga b. Sangerhausen, Germany. | Museum für Naturkunde, Berlin, Germany. |
| MB.Pb.1993/10027* (**Fig. 5D**) | Unpublished | Quaternary (2.58-0 Ma) | Seeberg, Germany. | Museum für Naturkunde, Berlin, Germany. |
| **CLUSTER I – not treated (bibliography only)** | | | | |
| Identified as *Solanum dulcamara* | Reid (1923) | Late Pliocene (3.6-2.58 Ma) | Pont-de-Gail, Cantal, France. | Rijksopsporing van Delfstoffen collection, The Netherlands. |
| *Tzeltalia calidaria* (Standl. & Steyerm.) E.Estrada & M.Martínez | Lundell 19625 (TEX) | Current (0 Ma) | Na | TEX |
| **CLUSTER J – PHYSALIS PLIOCENICA – not treated (bibliography only)** | | | | |
| Holotype *Physalis pliocenica* | Szafer (1946) | Pliocene (5.33 -2.58 Ma) | Kroscienko, Poland. | Na |
| 14183-28:3 – Identified as *Solanum nigrum* L. | van der Burgh (1987) | Late Miocene (11.63-5.33 Ma) | North Rhine, Hambach, Germany. | Laboratory of Paleobotany and Palynology, Utrecht, The Netherlands. |
| *Lycianthes nitida* Bitter | Dean (2020) | Current (0 Ma) | Na | Na |
| *Anisodus tanguticus* Pascher | Zhang & al. (2005) | Current (0 Ma) | Na | Na |
| **CLUSTER K – HYOSCYAMUS UNDULATUS** | | | | |
| K587/30 – Identified as *Solanum dulcamara* L. (**Fig. 5F**) | Dorofeev (1977) | Late Miocene / Early Pliocene (11.63-3.6 Ma) | 54.753123, 54.883264, Bashkiria, Ural, Russia. | Komarov Botanical Institute, St. Petersburg, Russia. |
| K453* – Identified as *Hyoscyamus niger* L. | Dorofeev (1957a, 1966) | Middle Pliocene (4.4-3.5 Ma) | 47.762692, 42.746863, Rostov oblast, Northern Caucasus, Russia. | Komarov Botanical Institute, St. Petersburg, Russia. |
| K345 – Identified as *Solanum nigrum* L. | Dorofeev (1956) | Pleistocene (2.6-0 Ma) | 48.484398, 44.786488, Volgograd oblast, Volga region, Russia. | Komarov Botanical Institute, St. Petersburg, Russia. |
| H135* | Unpublished | Na | 56.030961, 83.895555, Tomsk oblast, West Siberia, Russia. | Komarov Botanical Institute, St. Petersburg, Russia. |
| H1280* | Unpublished | Neogene (23.03-5.33 Ma) | 53.354407, 82.237467, Altay kray, West Siberia, Russia. | Komarov Botanical Institute, St. Petersburg, Russia. |
| K531* | Unpublished | Early Pliocene (5.33-3.6 Ma) | 42.828644, 41.504082, Duab, Abkhazia, Georgia. | Komarov Botanical Institute, St. Petersburg, Russia. |
| MB.Pb.1993/7754 | Unpublished | Pliocene (5.33 -2.58 Ma) | Kroscienko, Poland. | Museum für Naturkunde, Berlin, Germany. |
| MB.Pb.1993/6832 | Unpublished | Late Pliocene (3.6-2.58 Ma) | Nordhausen, Germany. | Museum für Naturkunde, Berlin, Germany. |
| K388 – Identified as *Physalis* sp. | Dorofeev (1963) | Miocene (23.03-5.33 Ma) | 56.343530, 84.087431, Tomsk oblast, West Siberia, Russia. | Komarov Botanical Institute, St. Petersburg, Russia. |
| H4008/93* **(Fig. 5G)** | Unpublished | Middle Miocene (15.97-11.63 Ma) | 53.997642, 76.373463, Pavlodar, Kazakhstan. | Komarov Botanical Institute, St. Petersburg, Russia. |
| H4895/49 – Holotype (designated here, **Fig. 5E**) | Unpublished | Early Miocene (23.03-15.97 Ma) | 55.814554, 84.524896, Novosibirsk oblast, West Siberia, Russia. | Komarov Botanical Institute, St. Petersburg, Russia. |
| K366* – Identified as *Solanum nigrum* L. | Dorofeev (1957b) | Late Miocene/ Early Pliocene (11.63-3.6 Ma) | 55.448199, 51.40682, Tataria, Volga region, Russia. | Komarov Botanical Institute, St. Petersburg, Russia. |
| H4390* | Unpublished | Early-middle Neopleistocene (2.58-0 Ma) | 57.495903, 73.844962, Omsk oblast, West Siberia, Russia. | Komarov Botanical Institute, St. Petersburg, Russia. |
| *Hyoscyamus reticulatus* L. | KIB3847 | Current (0 Ma) | Na | Komarov Botanical Institute, St. Petersburg, Russia. |
| *Hyoscyamus pallidus* Waldst. & Kit. ex Willd. | KIB7396a | Current (0 Ma) | Na | Komarov Botanical Institute, St. Petersburg, Russia. |
| *Hyoscyamus muticus* L. | KIB7396b | Current (0 Ma) | Na | Komarov Botanical Institute, St. Petersburg, Russia. |
| *Hyoscyamus albus* L. | KIB7396c | Current (0 Ma) | Na | Komarov Botanical Institute, St. Petersburg, Russia. |
| *Lycianthes multiflora* Bitter (**Fig. 5H)** | BM001071257 | Current (0 Ma) | Na | BM |
| **CLUSTER L - SOLANOIDES DOROFEEVII** | | | | |
| H4220 | unpublished | Late Pliocene (3.6-2.58 Ma) | 57.131842, 74.603660, Omsk oblast, West Siberia, Russia. | Komarov Botanical Institute, St. Petersburg, Russia. |
| H4736 | unpublished | Early Miocene (23.03-15.97 Ma) | 54.710437, 77.821560, Novosibirsk oblast, Russia. | Komarov Botanical Institute, St. Petersburg, Russia. |
| Identified as *Hyoscyamus niger* L. | Reid (1923) | Late Pliocene (3.6-2.58 Ma) | Pont-de-Gail, Cantal, France. | Rijksopsporing van Delfstoffen collection, The Netherlands. |
| Identified as *Physalis pliocenica* | Szafer (1961) | Miocene: Tortonian (11.63-7.25 Ma) | Stare Gliwice, Poland. | -- |
| K530/21* – Holotype (designated here) – Identified as *Physalis* sp. – *Solanum* sp. **(Fig. 6A)** | Dorofeev (1964) | Middle Miocene (15.97-11.63 Ma) | 44.515202, 39.730509, Krasnodar kray, Northern Caucasus, Russia. | Komarov Botanical Institute, St. Petersburg, Russia. |
| RGM793435 – Identified as *Physalis alkekengi* L. (currently *Alkekengi officinarum* Moench) | Reid & Reid (1910) | Pliocene (5.33 -2.58 Ma) | Tagelen, Limburg, The Netherlands. | Naturalis Biodiversity Center, Leiden, The Netherlands. |
| MRSN-P/345-CCN4334 | Martinetto & Festa (2013) | Pleistocene: Gelasian (2.58-1.8 Ma) | Buronzo, Italy. | CENOFITA collection, managed by the Regional Museum of Natural Sciences of Turin, Turin, Italy. |
| K432 – *Solanum dulcamara* L. | Dorofeev (1951, 1955) | Middle-Late Miocene (15.97-5.33 Ma) | 46.410147, 30.758181, Odesa, Odesa oblast, Ukraine. | Komarov Botanical Institute, St. Petersburg, Russia. |
| H4148 | unpublished | Early Oligocene (33.9-27.82 Ma) | 60.108504, 83.242754, Tomsk oblast, West Siberia, Russia. | Komarov Botanical Institute, St. Petersburg, Russia. |
| K266 – *Physalis* sp. | Dorofeev (1988) | Middle Miocene (15.97-11.63 Ma) | 51.91591, 42.469724, Tambov oblast, Central Black Earth region, Russia. | Komarov Botanical Institute, St. Petersburg, Russia. |
| H2727 | unpublished | Late Oligocene / Early Miocene (27.82-15.97 Ma) | 56.241528, 77.358989, Novosisibirsk oblast, West Siberia, Russia | Komarov Botanical Institute, St. Petersburg, Russia. |
| H4000 | unpublished | Late Oligocene (27.82-23.03 Ma) | 54.020703, 77.512922, Novosibirsk oblast, West Siberia, Russia. | Komarov Botanical Institute, St. Petersburg, Russia. |
| H2993 | unpublished | Early Miocene (23.03-15.97 Ma) | 64.836188, 82.933646, Yamalo-Nenets autonomous okrug, West Siberia, Russia. | Komarov Botanical Institute, St. Petersburg, Russia. |
| K518 | unpublished | Oligocene (33.9-23.03 Ma) | 60.100319, 83.209543, Tomsk oblast, West Siberia, Russia. | Komarov Botanical Institute, St. Petersburg, Russia. |
| H1341* | unpublished | Na | 61.425672, 82.885153, Khanty-Mansy autonomous district, Tyumen oblast, West Siberia, Russia. | Komarov Botanical Institute, St. Petersburg, Russia. |
| H4008 | unpublished | Middle Miocene (15.97-11.63 Ma) | 53.997642, 76.373463, Pavlodar, Kazakhstan. | Komarov Botanical Institute, St. Petersburg, Russia. |
| MB.Pb.1993/8849 | unpublished | Middle Miocene (15.97-11.63 Ma) | Berzdorf, Germany. | Museum für Naturkunde, Berlin, Germany. |
| MB.Pb.1993/7753* | unpublished | Pliocene (5.33-2.58 Ma) | Kroscienko, Poland. | Museum für Naturkunde, Berlin, Germany. |
| MB.Pb.1993/5546B **(Fig. 6C)** – Identified as *Physalis* | Mai & Walther (1988) | Late Pliocene (3.6-2.58 Ma) | Berga b. Sangerhausen, Germany. | Museum für Naturkunde, Berlin, Germany. |
| MB.Pb.1993/5545 – Identified as *Scopolia carniolica* Jacq. | Mai & Walther (1988) | Late Pliocene (3.6-2.58 Ma) | Berga b. Sangerhausen, Germany. | Museum für Naturkunde, Berlin, Germany. |
| MB.Pb.1993/11406 | unpublished | Late Pliocene (3.6-2.58 Ma) | Öbel, The Netherlands. | Museum für Naturkunde, Berlin, Germany. |
| MB.Pb.1993/6802* (**Fig. 1D**) | unpublished | Pleistocene: Gelasian (2.58-1.8 Ma) | Rusell-Tiglia Egypte / Tegelen, The Netherlands. | Museum für Naturkunde, Berlin, Germany. |
| MRSN-P/345-CCN3059a | Cavallo & Martinetto (2001) | Pleistocene: Gelasian (2.58-1.8 Ma) | Castelletto Cervo II, Italy. | CENOFITA collection, managed by the Regional Museum of Natural Sciences of Turin, Turin, Italy. |
| MRSN-P/345-CCN1966B | Cavallo & Martinetto (2001) | Pleistocene: Gelasian (2.58-1.8 Ma) | Castelletto Cervo II, Italy. | CENOFITA collection, managed by the Regional Museum of Natural Sciences of Turin, Turin, Italy. |
| MRSN-P/345-CCN3059b | Cavallo & Martinetto (2001) | Pleistocene: Gelasian (2.58-1.8 Ma) | Castelletto Cervo II, Italy. | CENOFITA collection, managed by the Regional Museum of Natural Sciences of Turin, Turin, Italy. |
| MRSN-P/345-CCN4324* **(Fig. 6B)** | Niccolini & al. (2022) | Late Miocene (5.6-5.33 Ma) | Ciabot Cagna, Corneliano d'Alba, Italy. | CENOFITA collection, managed by the Regional Museum of Natural Sciences of Turin, Turin, Italy. |
| MRSN-P/345-CCN4321* | unpublished | Pleistocene: Calabrian (1.8-0.77 Ma) | Arda, Italy. | CENOFITA collection, managed by the Regional Museum of Natural Sciences of Turin, Turin, Italy. |
| *Lycianthes pringlei* Bitter | Dean (2007) | Current (0 Ma) | Na | Na |
| *Lycianthes surotatensis* Gentry | Dean (2017) | Current (0 Ma) | Na | Na |
| *Lycianthes jaliscensis* E. Dean | Dean (2017) | Current (0 Ma) | Na | Na |
| *Lycianthes heteroclita* Bitter | Dean (2020) | Current (0 Ma) | Na | Na |
| *Lycianthes geminiflora* Bitter | Dean (2020) | Current (0 Ma) | Na | Na |
| *Capsicophysalis potosina* (B.L.Rob. & Greenm.) Averett & M.Martínez | BM000775314 | Current (0 Ma) | Na | BM |
| *Deprea pauciflora* Deanna, Barboza & S.Leiva | Deanna & al. (2016) | Current (0 Ma) | Ecuador. | Na |
| *Deprea macasiana* (Deanna, S.Leiva & Barboza) Barboza | Deanna & al. (2014) | Current (0 Ma) | Ecuador. | Na |
| *Saracha nigribaccata* J.M.H.Shaw | Smith 212 (COLO) | Current (0 Ma) | Na | COLO |
| *Exodeconus miersii* (Hook.f.) D'Arcy | Schmitt 108 (NY) | Current (0 Ma) | Na | NY |
| *Iochroma cyaneum* (Lindl.) M.L.Green | Smith 265 (COLO) | Current (0 Ma) | Na | COLO |
| *Deprea hawkesii* (Hunz.) Deanna | Deanna & al. 438 (CORD) | Current (0 Ma) | Na | CORD |
| *Darcyanthus spruceanus* (Hunz.) Hunz. | Barboza s.n. (CORD) | Current (0 Ma) | Na | CORD |
| *Benthamiella* sp. | Donat 191 (NY) | Current (0 Ma) | Na | NY |
| *Archiphysalis chamaesarachoides* (Makino) Kuang | K. Chiba s.n. (CORD00028525) | Current (0 Ma) | Na | CORD |
| *Physaliastrum heterophyllum* (Hemsl.) Migo | Deng & Yan 79035 (NY) | Current (0 Ma) | Na | NY |
| *Nolana spathulata* (Ruiz & Pav.) Mesa | FLSP 1044 (NY) | Current (0 Ma) | Na | NY |
| *Deprea cardenasiana* Hunz. | Deanna 426 (CORD) | Current (0 Ma) | Perú. | CORD |
| *Brachistus stramonifolius* Miers | Bye 3561 (COLO) | Current (0 Ma) | Na | COLO |
| *Witheringia stellata* (Greenm.) Hunz. | Hernandez Magana 5327 (COLO) | Current (0 Ma) | Na | COLO |
| *Vassobia breviflora* (Sendt.) Hunz. | Martínez 1059 (CORD00097043) | Current (0 Ma) | Na | CORD |
| *Lycianthes parasitica* Bitter | E00426956 | Current (0 Ma) | Na | E |
| *Vestia lycioides* Willd. | West 4986 (NY) | Current (0 Ma) | Na | NY |
| *Calliphysalis carpenteri* (Riddell) Whitson **(Fig. 6D)** | Ward 5893 (NY) | Current (0 Ma) | Na | NY |
| **CLUSTER M – ALBIONITES ARNENSIS** | | | | |
| V40898 – *Albionites arnensis* **(holotype, Fig. 6E)** | Chandler (1962) | Eocene: late Ypresian / early Lutetian (48-46 Ma) | Arne, Poole Formation, United Kingdom. | Natural History Museum, London, United Kingdom. |
| *Lycianthes oliveriana* (Lauterb. & Schum.) Bitter **(Fig. 6F)** | Hartley 10136 (A) | Current (0 Ma) | Na | A |
| **CLUSTER N – not treated (bibliography only)** | | | | |
| Identified as *Solanum dulcamara* L. | Szafer (1946) | Pliocene (5.33 -2.58 Ma) | Kroscienko, Poland. | -- |
| **CLUSTER O – NEPHROSEMEN RETICULATUM** | | | | |
| UF6500 – *Nephrosemen reticulatum* – Holotype (**Fig. 6G**) | Manchester (1994) | Middle Eocene (47.8-37.7 Ma) | Nut Beds flora, Clarno Formation, north-central Oregon, United States. | Florida Museum of Natural History, University of Florida, United States. |
| UF9380 – identified as *Nephrosemen reticulatum* | Manchester (1994) | Middle Eocene (47.8-37.7 Ma) | Nut Beds flora, Clarno Formation, north-central Oregon, United States. | Florida Museum of Natural History, University of Florida, United States. |
| UF9378 – identified as *Nephrosemen reticulatum* | Manchester (1994) | Middle Eocene (47.8-37.7 Ma) | Nut Beds flora, Clarno Formation, north-central Oregon, United States. | Florida Museum of Natural History, University of Florida, United States. |
| UF9777 – identified as *Nephrosemen reticulatum* | Manchester (1994) | Middle Eocene (47.8-37.7 Ma) | Nut Beds flora, Clarno Formation, north-central Oregon, United States. | Florida Museum of Natural History, University of Florida, United States. |
| UF9382 – identified as *Nephrosemen reticulatum* | Manchester (1994) | Middle Eocene (47.8-37.7 Ma) | Nut Beds flora, Clarno Formation, north-central Oregon, United States. | Florida Museum of Natural History, University of Florida, United States. |
| UF9376– identified as *Nephrosemen reticulatum* | Manchester (1994) | Middle Eocene (47.8-37.7 Ma) | Nut Beds flora, Clarno Formation, north-central Oregon, United States. | Florida Museum of Natural History, University of Florida, United States. |
| UF9381 – identified as *Nephrosemen reticulatum* | Manchester (1994) | Middle Eocene (47.8-37.7 Ma) | Nut Beds flora, Clarno Formation, north-central Oregon, United States. | Florida Museum of Natural History, University of Florida, United States. |
| MB.Pb.2000/16 – identified as *Hyoscyamus reticulatus* L. | Mai & Velitzelos (2007) | Pliocene / Pleistocene (5.33-0.01 Ma) | Kallithea, Rhodes, Greece. | Museum für Naturkunde, Berlin, Germany. |
| *Trozelia umbellata* Raf. **(Fig. 5H)** | Särkinen & al. 4783 (E) | Current (0 Ma) | Na | E |
| **CLUSTER P – CAPSICUM PLIOCENICUM** | | | | |
| MRSN-P/345-CCN6065b* – Holotype (designated here; **Fig. 7A**) | Unpublished | Pleistocene (0.78-0.13 Ma) | Bucine, Italy. | CENOFITA collection, managed by the Regional Museum of Natural Sciences of Turin, Turin, Italy. |
| RGM792900 – Identified as *Solanum* | Reid & Reid (1915) | Pliocene (5.33 -2.58 Ma) | Reuver, Limburg, The Netherlands. | Naturalis Biodiversity Center, Leiden, The Netherlands. |
| RGM793332 – Identified as *Solanum dulcamara* L. | Reid & Reid (1907) | Pliocene (5.33 -2.58 Ma) | Tegelen, Limburg, The Netherlands. | Naturalis Biodiversity Center, Leiden, The Netherlands. |
| Identified as *Solanum dulcamara* L. | Reid & Reid (1907) | Pliocene (5.33 -2.58 Ma) | Tegelen-Sur-Meuse, Limburg, The Netherlands. | Rijksopsporing van Delfstoffen collection, The Netherlands. |
| Identified as *Solanum* sp. | Reid & Reid (1915) | Pliocene (5.33 -2.58 Ma) | Limburg and Prussian border, The Netherlands. | Rijksopsporing van Delfstoffen collection, The Netherlands. |
| *Capsicum baccatum* L. **(Fig. 7B)** | Barboza & al. (2022) | Current (0 Ma) | Na | Na |
| **CLUSTER Q – THANATOSPERMA MINUTUM** | | | | |
| MB.Pb.2003/0073 – Holotype (designated here) – identified as *Solanum dulcamara* L. **(Fig. 7C)** | Gümbel & Mai (2004) | Late Pliocene (3.6-2.58 Ma) | Oberzella a. d. Werra, Germany. | Museum für Naturkunde, Berlin, Germany. |
| MRSN-P/345-CCN448 – cf. *Hyoscyamus* | Cavallo & Martinetto (2001) | Pleistocene: Gelasian (2.58-1.8 Ma) | Castelletto Cervo II, Italy. | CENOFITA collection, managed by the Regional Museum of Natural Sciences of Turin, Turin, Italy. |
| MRSN-P/345-CCN3067 | Girotti & al. (2003) | Pleistocene: Gelasian (2.58-1.8 Ma) | Torre Picchio, Italy. | CENOFITA collection, managed by the Regional Museum of Natural Sciences of Turin, Turin, Italy. |
| H676/80a* **(Fig. 7D)** | Unpublished | Holocene (0.01-0.001 Ma) | 55.166345, 82.857844, Novosibirsk oblast, West Siberia, Russia. | Komarov Botanical Institute, St. Petersburg, Russia. |
