## Supplementary table S2 for "Seed fossil record of Solanaceae revisited"

**Table S2.** Seed traits scored for fossil specimens and extant species of Solanaceae. **Bold** characters are secondary traits, calculated from primary measurements; *italicized* characters were not included in the NMDS analysis. Characters marked with an asterisk (*) were scored using high-resolution micro-CT. Roundness compares the shape of an object to a perfect circle, with a value of 1 indicating a perfect circle and values closer to 0 representing more elongated or irregular shapes.

| Seed character | # | Continuous (C) / Discrete (D) | Character states |
| --- | --- | --- | --- |
| Length | 1 | C | -- |
| *Width* | 2 | C | -- |
| **Length/width ratio** | 3 | C | -- |
| Area | 4 | C | -- |
| Compression | 5 | D | 0= flattened  1= not flattened |
| Hilum position | 6 | D | 0= lateral or sub-lateral  1= terminal |
| Embryo shape* | 7 | D | 0 = curved  1= straight  2 = circinate/coiled |
| Hilar-chalazal cavity | 8 | D | 0= absent  1= present |
| Exotestal cell walls | 9 | D | 0= straight  1= sinuate-cerebelloid |
| Wings | 10 | D | 0= absent  1= present |
| Elaiosomes* | 11 | D | 0= absent  1= present |
| Exotestalcells area | 12 | C | -- |
| Exotestal cells perimeter | 13 | C | -- |
| Exotestal **cells perimeter/area ratio** | 14 | C | -- |
| Exotestal cells roundness | 15 | C | -- |
