## Supplementary table S6 for "Seed fossil record of Solanaceae revisited"

**Table S6.** Discrete and continuous characters analyzed, including scores that show the contribution of each variable to each axis of the NMDS and significance. The coding scheme for each character is shown in Table S2. Significance codes: 0 ‘***’ 0.001 ‘**’ 0.01 ‘*’ 0.05 ‘.’ 0.1 ‘ ’ 1.

|  | NMDS1 | NMDS2 | r2 | Pr(>r) | Significance |
| --- | --- | --- | --- | --- | --- |
| length | -0.02342 | -0.99973 | 0.0751 | 0.002 | ** |
| length/width ratio | 0.87109 | -0.49113 | 0.0938 | 0.001 | *** |
| area | 0.03707 | -0.99931 | 0.0527 | 0.017 | * |
| compression | 0.98323 | -0.18235 | 0.6955 | 0.001 | *** |
| hilum position | 0.23277 | 0.97253 | 0.2152 | 0.001 | *** |
| embryo shape | -0.32267 | -0.94651 | 0.1446 | 0.001 | *** |
| hilar-chalazal cavity | -0.82415 | -0.56636 | 0.731 | 0.001 | *** |
| testal cell walls | -0.42116 | 0.90699 | 0.646 | 0.001 | *** |
| wings | -0.6267 | -0.77926 | 0.0649 | 0.005 | ** |
| elaiosomes | -0.93704 | 0.34921 | 0.0935 | 0.001 | *** |
| testal cells area | 0.42497 | -0.90521 | 0.0633 | 0.002 | ** |
| testal cell perimeter | 0.21445 | -0.97673 | 0.0577 | 0.008 | ** |
| testal cells perimeter/area ratio | 0.2577 | 0.96622 | 0.0375 | 0.058 | . |
| testal cells roundness | -0.08737 | -0.99618 | 0.0107 | 0.484 |  |
